# Micro- to nano-scale investigation of Precambrian metasediments: biogenicity and preservation in the 3.22 Ga Moodies Group (Barberton Greenstone Belt, S. Africa) and the 2.46 Ga Brockman Iron Formation (Hamersley Basin, W. Australia)

**DOI:** 10.1101/2023.02.08.527618

**Authors:** Hervé Bellon, Jacek Gieraltowski, François Michaud, Gaëlle Simon, Stéphane Cerantola, Martin Homann, Ian Foster, Pascal Ballet, Stefan V. Lalonde

## Abstract

Precambrian metasediments provide a unique archive for understanding Earth’s earliest biosphere, however traces of microbial life preserved in ancient rocks are often controversial. In this study we leveraged several micro- to nano-scale techniques to study filamentous structures previously reported in clastic sediments of the 3.22 Ga Moodies Group, Barberton Greenstone Belt, S. Africa. We performed petrographic, mineralogical, electron microprobe, confocal fluorescence and electron microscopy analyses of these structures in order to evaluate their biogenicity and syngenecity. We also examined drill core samples of deep-water iron formations from the 2.46 Ga Joffre member of the Brockman Iron Formation (Hamersley Basin, W. Australia) to better understand their potential biogenicity. In both cases, we aimed to resolve primary vs. secondary mineral assemblages and their relation to filamentous or sedimentary structures. In the Moodies Group samples, filamentous structures were resolved by confocal imaging and revealed to be crosscut by later metamorphic phases, highlighting their syngenetic nature. Three-dimensional imaging reveals that while the filamentous structures are not necessarily associated with grain boundaries (e.g., as organic coatings), they form both sheets and filaments, complicating their interpretation but not ruling out a biological origin. No organic microstructures appeared to be preserved in our Dales Gorge samples. We also examined the possible application of electron paramagnetic resonance spectroscopy (EPR) to carbonaceous matter in ancient silica-rich matrices, similar to Bourbin et al. (2013), using samples from the Brockman iron formation. While resonance associated with organic matter was largely unresolvable in the Brockman iron formation samples due to their low organic matter contents, large effects on the EPR spectra were apparent stemming from the presence of magnetic iron minerals, highlighting the need to carefully consider sample composition in EPR analyses targeting ancient organic matter. Collectively, this study highlights the added value of micro- to nano-scale techniques as applied to Precambrian metasediments containing traces of ancient life, for example in revealing the pre-metamorphic emplacement and three-dimensional structure of filaments in the Moodies Group, but also the potential drawbacks and pitfalls, such as the case of strong magnetic mineral interference in EPR analysis of organic matter in trace abundance in the Dales Gorge.

## 1. Introduction

The vast majority of Earth’s most ancient purported microfossils are hosted in microcrystalline silica minerals in chert. The organic matter constituting these fossils is generally highly mature and graphitized, such that organic biomarkers provide little information as to biogenicity, and other means of verification are required. Other evidence used to demonstrate biogenicity and syngenicity includes geological context, microfossil morphology, relationships between microfossils and surrounding minerals, and evidence against secondary emplacement. The interpretation of ancient filamentous or coccoid structures as microfossils has proven to be highly contentious. For example, in the 3.46 Ga Apex Chert (W. Australia), filamentous structures purported to represent fossilized bacteria, including putative cyanobacteria (Schopf 1993), were later dismissed as artifacts associated with hydrothermal emplacement of kerogen of potentially abiotic origin (Brasier et al. 2002).

More recently, filamentous microstructures have been reported in iron- and silica-rich chemical sediments from the >3.77 Ga Nuvvuagittuq belt in Quebec, Canada (Dodd et al. 2017). The microstructures reported in this work may constitute the oldest available evidence for life on Earth, and accordingly, extensive efforts were made to establish biogenicity. It is perhaps not surprising that filamentous bacteria may be preserved in iron-rich chemical sediments, as modern analogues, such as the iron-oxidizing bacterial mats found growing at the Arctic Mid-Ocean Ridge (Johannessen et al. 2017) or on the Lohihi seamount offshore of Hawaii (Karl et al. 1989), show abundant filamentous structures preserved throughout the mats.

However, in the rock record, besides the filamentous microstructures reported in the Dodd et al. (2017) study, such structures are generally rare in iron-rich cherty matrices. One of the most studied examples is in the Gunflint chert, where micro-digitate stromatolites containing filamentous microstructures are found to be encrusted in hematite. The formation of these microstructures is controversial. While their biogenicity is generally unquestioned, there exist significant doubts surround the syngenetic emplacement of the iron oxide minerals; Shapiro and Konhauser (2015) suggested that rather than representing a primary feature, Fe-encrustation occurred later during fluid flow through the deposit post-burial.

Some chert-rich rocks that were deposited throughout the Precambrian are conspicuously free of microfossils, notably Precambrian Banded Iron Formations (BIF). BIF are iron- and silica-rich chemical sediments (>15% Fe, Gross 1965) that were deposited throughout much of the Precambrian. While these sediments are highly siliceous and are generally considered to have been deposited as the result of microbial activity (see Konhauser et al. 2017, for review), they generally lack abundant microfossils, contrary to their pure-silica counterparts, chemical cherts.

Precambrian microfossils are not only restricted to cherty metasediments. In the 3.22 Ga Moodies Group (Barberton Greenstone Belt, S. Africa), purported microbial mats preserved in subtidal to intertidal sandstones contained filamentous microstructures of probable biological origin. These microbial mat communities have been previously described by Homann et al. (2015), who characterized them using light microscopy and μXRF imaging, however little work has been performed on the filamentous structures hosted within.

Over the last few decades, emerging micro- and nano-scale analytical techniques have reinforced our capacity for recognizing biological vs. abiogenic structures in ancient rocks. For example, instruments capable of spatially resolved X-Ray Diffraction independent of electron microscope instruments permit rapid analyses of mineral distributions at the micron-scale. The proliferation of electron microprobe instruments now permits such analyses to be coupled with in-situ chemical analyses at high precision and ultra-high spatial resolution (routinely down to 1 micron). Confocal laser scanning microscopy permits optical and fluorescence imaging at high spatial resolution at multiple depths within samples, and via image stacking, permits “optical sectioning” of samples in thin section to provide a three-dimensional view of rock-hosted microstructures. Furthermore, excitation with lasers of different wavelengths, combined with bandpass filters permitting imaging at different fluorescence wavelengths, permits materials of different composition and fluorescence properties to be resolved.

Electron paramagnetic resonance (EPR) analysis is an additional technique with proven application to Precambrian rocks of purported biological origin but has received relatively little attention compared to the techniques mentioned above (Skrzypezak-Bonduelle et al. 2008; Bourbin et al. 2013; Gourier et al. 2019). Briefly, EPR is a spectroscopic technique that is approximately 1000 times more sensitive than nuclear magnetic resonance imaging that examines paramagnetic properties of natural and artificial samples. It is based on the adsorption of electromagnetic emission by atoms containing one or more unpaired electrons, whereby under the influence of a magnetic field, their electron spin is broken into two components of opposite spin by the so-called Zeeman effect. At excitation energies in the X-ray range, such electrons enter into resonance, with characteristic adsorption energies for different nuclei that are expressed as a function of excitation energy. Organic radicals respond to this technique, and furthermore, their response evolves as a function of thermal maturity. This property has been used to evaluate the syngenecity of ancient organic matter, and even to propose potential age dating schemes based on properties of the EPR spectra of ancient organic matter (Skrzypezak-Bonduelle et al. 2008; Bourbin et al. 2013).

We applied all of the above techniques to better understand primary vs. secondary mineral assemblages, as well as their relation to filamentous microstructures when present, in samples from two Precambrian sedimentary units of purported strong biological influence: microbial-mat-bearing sandstones of the 3.22 Ga Moodies Group, and iron- and silica-rich chemical metasediments from the 2.46 Ga Joffre Member of the Brockman Iron Formation.

## 2 Geological Overview

### 2.1 Iron formations and microbial mats of the Moodies Group

The up ~3 km thick siliciclastic Moodies Group is the uppermost unit of the Barberton Greenstone Belt (3.55 to ca. 3.22 Ga) of South Africa and Swaziland that is located at the eastern margin of the Kaapvaal Craton (Lowe and Byerly 1999). The sandstones of the Moodies Group are underlain by the volcaniclastic Fig Tree Group and the volcanic-dominated Onverwacht Group, respectively (Fig. 1). Depositional ages of volcanic beds throughout the Moodies Group, obtained through single-zircon U–Pb dating, indicate that deposition began ~3223±1 Ma and had ended by ~3219 ± 9 Ma (De Ronde and Kamo 2000; Heubeck et al. 2013). Consequently, the Moodies succession was deposited and deformed within <1–14 Ma and thus represents a unique, high-resolution archive of Archean surface and sedimentation processes. The Moodies Group is also well known for the occurrence of locally abundant, macroscopically visible carbonaceous laminations interpreted as fossil remains of microbial mats (Noffke et al. 2006; Heubeck 2009; Homann et al. 2015). Raman microspectroscopy demonstrates that these laminations experienced peak temperatures of ~365°C (Tice et al. 2004; Homann et al. 2018), which is consistent with the metamorphic grade of the Moodies Group determined by chlorite Al^IV^ geothermometry (Xie et al. 1997) and thus proves the syngenetic origin of the mats.

**Figure 1.**
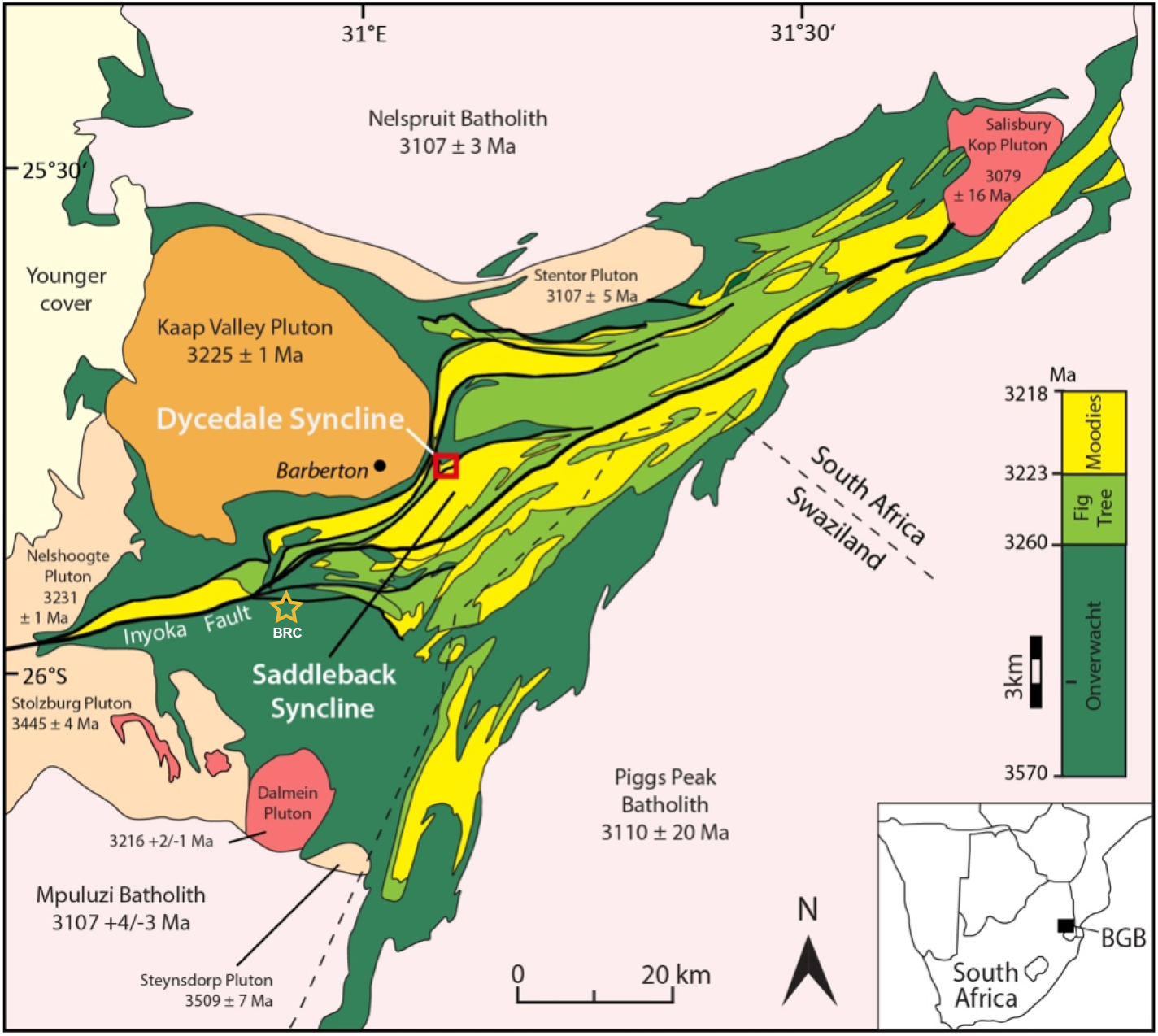
Map of the Barberton Greenstone Belt showing the major sedimentary groups as well as the location of the Dycedale syncline within the Moodies Group. The orange star labeled “BRC” represents the locality for the Buck Reef Chert sample used in EPR mixing tests. Figure reproduced from Homann et al. (2018).

### 2.2 Dales Gorge and Joffre members of the Brockman Iron Formation

The Brockman Iron Formation of the Hamersley Group, Western Australia, is found on the Pilbara craton and is comprised approximately 600 m of chemical sediments (iron formation) and shales (Fig. 2). From bottom to top, it consists of the ~140 m thick Dales Gorge Member (iron formation), the ~13m thick Whaleback Shale, the ~360 m Joffre Member (iron formation), and is capped by the ~50 m thick Yandicoogina Shale. It is constrained in age by a 2495 ± 3 Ma date at the base of the Dales Gorge member and dates of 2459 ± 3 and 2454 ± 3 Ma in the Joffre member, all determined by SHRIMP U-Pb dating of zircon grains (Pickard 2002; Trendall et al. 2004). Deposition rates for the Dales Gorge member are estimated at 5 m/Myr, and 180 m/Myr for the Joffre member (Trendall et al. 2004). Microbanding is well known in this deposit, plausibly representing annual varves (Trendall and Blockley 1970). The Brockman Iron Formation is not thought to have experienced burial metamorphism temperatures in excess of 160° C (Trendall 1966). This low grade of metamorphism is exceptional for this age and favors the preservation of mineral assemblages and nano-scale textures reflecting early diagenetic and metamorphic BIF mineralogical transformations.

**Figure 2.**
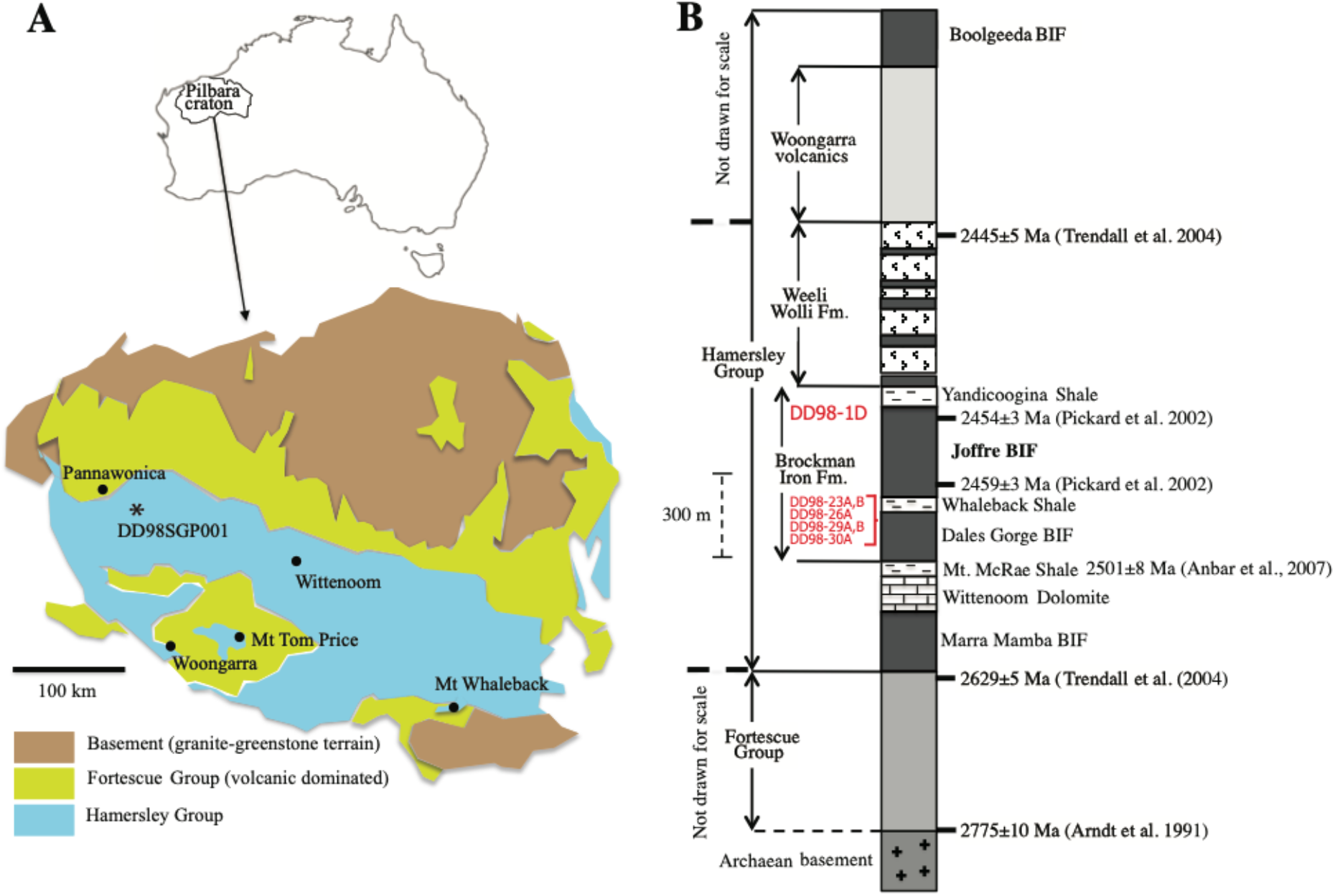
(A) Situation of the drill core DD98SGP001 in the Hamersley basin of the Pilbara craton, NW Australia. (B) Stratigraphic column of the Hamersley Group with sample locations indicated with red text. Figure adapted from Haugaard et al. (2015).

## 3. Sampling and Methods

### Sampling and sample preparation

Samples from the Moodies Group were contributed by Martin Homann, collected from field exposures between 2011 and 2013 from the Saddleback Syncline in the Barberton Greenstone Belt.

Samples from the Joffre Member were contributed by Kurt Konhauser and Stefan Lalonde, sampled from drill core DD98SGP001 at Rio Tinto plc in Perth. See Krapez et al. (2003) for a detailed description of this core. Powders were prepared using a tungsten carbine hammer crusher and powdered in an agate mill at the European Institute for Marine Studies, Université de Bretagne Occidentale, France. Thin sections were prepared at the University of Alberta Department of Earth and Atmospheric Sciences (Joffre member) or at Freie Universität Berlin (Moodies Group).

### EMPA Microprobe Analysis

Polished thins sections from both the Moodies Group (11-452-B5) and Brockman Formation (DD98-21 and 26A) were analyzed using a Cameca SX100 at IFREMER, Brest, France operating at 15kV and 20nA with a 1 μm spot size.

### Scanning Electron Microscopy

Powders (10 to 100 μm in size) as well as the polished thin section 11-452-B5 were analyzed using the HITACHI S-3200N microscope at the Plateforme d’Imagerie et de Mesures en Microscopie (PIMM) of the Université de Bretagne Occidentale. Semi-quantitative energy-dispersive x-ray spectrometry analyses were performed with an associated PGT Prism 2000 Si(Li) detector with counting times of 400 sec.

### Electron Paramagnetic Resonance Analyses

Five samples from the Brockman Iron Formation (DD98-1D, DD98-23A, DD98-23B, DD98-29A and DD98-29B with their respective locations in the stratigraphic column in Fig. 2) were selected for electron paramagnetic resonance (EPR) analyses on whole rock sample powders without later separation of the different phases. 5 to 10 mg of each powder was deposited in a quartz tube and analyzed using a Brucker Elesxys 500 EPR instrument operating at 9.3 GHz. The scanning frequency was varied in steps of 10 gauss from 0 to 10,000 gauss for the large-scale scans, and for the fine scale scans, over a corresponding range of 400 gauss centered around 3600 gauss in steps of 0.05 gauss.

### X Ray Diffraction

X-ray diffractograms of individual beds from two samples of the Brockman Iron Formation (samples DD98-26A and DD98-30A; see Figs. 2 and 5) were acquired using a PANalytical Empyrean X-Ray Diffractometer using Cu Kα1/Kα2 radiation, a 0.1 × 4 mm collimated beam, and with a PIXcel3D linear position-sensitive detector. Diffraction spectra were analysed using the X’Pert Highscore software and the associated crystallographic database PAN-ICSD (Inorganic Crystal Structure Database).

### Confocal Laser Scanning Microscopy

Confocal Laser Scanning Microscopy was performed at the Plateforme d’Imagerie et de Mesures en Microscopie (PIMM) of the Université de Bretagne Occidentale using a Zeiss confocal microscope equipped with 8 lasers spanning 405 to 635 nm and a GaSP spectral detector. For this study, imaging was performed using two excitation wavelengths simultaneously, 488 nm and 633 nm, with laser power at approximately 20-25%.

### Micro X-ray Fluorescence

Micro x-ray fluorescence (μXRF) was performed at the University of Brest using a Bruker M4 Tornado operating with an excitation energy of 50 kV with the source at 600 μA. Three different spot analyses were performed per sample and averaged to obtain representative concentrations. Concentration calibration was performed using the Fundamental Parameters algorithm built into the M4 Tornado software package.

## 4. Results and Discussion

### 4.1 Optical, electron microscope, and confocal imaging evidence for primary vs. secondary processes affecting Moodies Group kerogen

Kerogen believed to represent microbial mats remnants in Moodies Group coarse-grained sandstones were imaged by visible light microscopy, by transmission electron microscopy (TEM), and by laser confocal microscopy (Fig. 3). In visible light micrographs, areas of concentrated kerogen <1 mm in thickness are clearly seen lying bed-parallel and appear generally laterally continuous (Fig. 3A) and are associated with nano-crystalline clays resolvable only by TEM (Fig. 3B). Confocal imaging of a contact between kerogen-rich and kerogen-poor sandstone further reveals a complex mixture of mineral grains and amorphous kerogen. While most of the signal captured appears to correspond to true fluorescence, it’s important to note that some cases of reflection are also visible (e.g. strong signals in the green wavelength associated with a fissure at the bottom of Fig. 3C). The sandstone contains a mixture of highly angular grains, sub-rounded grains, and some well-rounded grains (Fig. 3C), and is poorly sorted. In places, individual grains are floating in areas of concentrated kerogen, while in other areas, kerogen is found squeezed between boundaries of grains that are in close contact (Fig. 3D).

**Figure 3.**
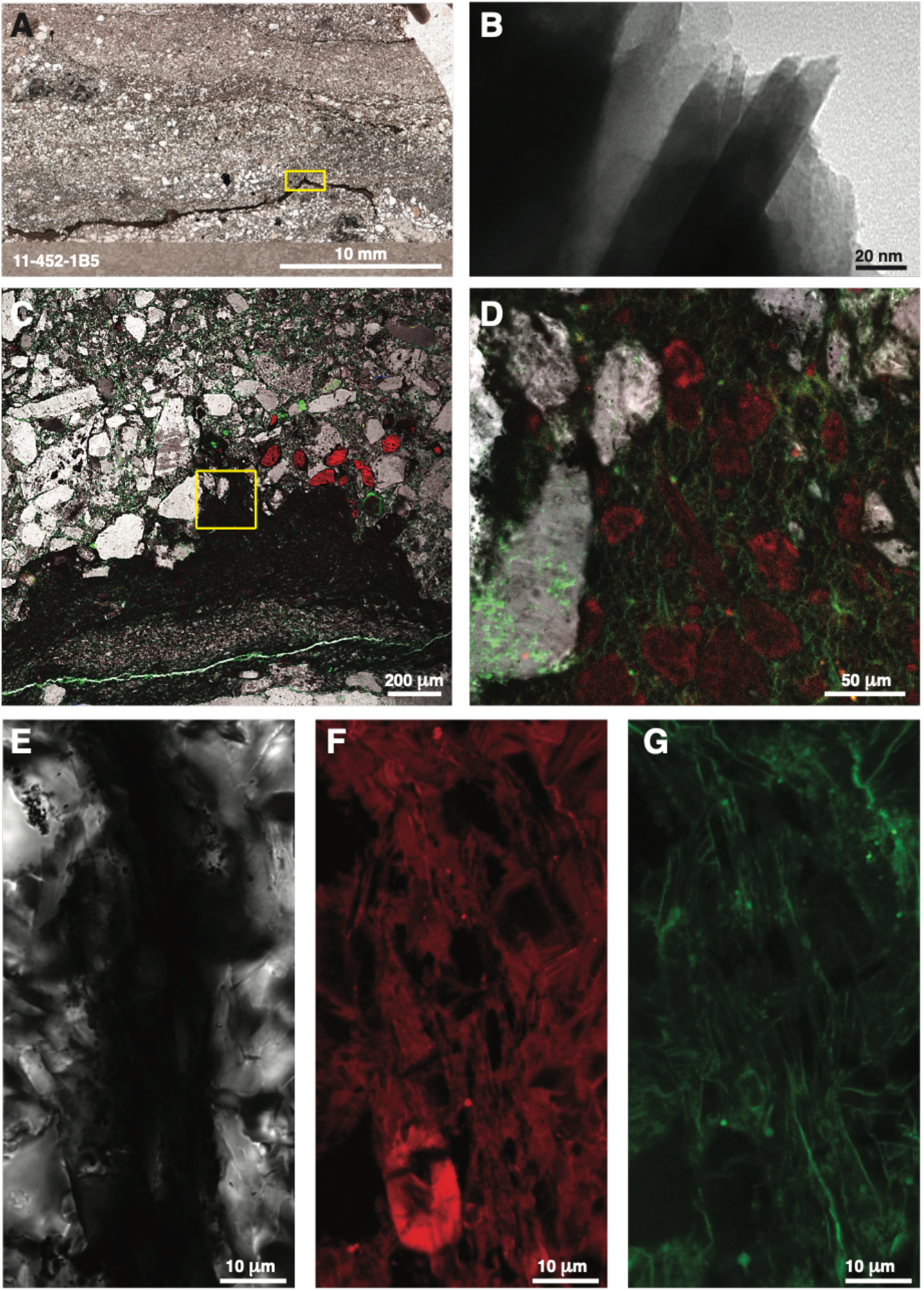
(A) Thin section 11-452-1B5 of microbial mats preserved in coarse-grained sandstone from the Dycedale syncline, 3.22 Ga Moodies Group, Barberton Greenstone Belt (S. Africa). (B) TEM image of nano-crystalline clays in the same thin section (C) Confocal Laser Scanning Microscope (CLSM) images of a flame structure (Flame Structure 1) in the same thin section (yellow box in (A)) showing white quartz crystals in visible light, secondary minerals and kerogen in red (633 nm excitation), and kerogen in green (488 nm excitation). (D) magnified image of yellow box in (C) showing secondary minerals (e.g., tourmaline) in red that crosscut filamentous structures imaged in green. (E, F, G) Quartz crystals in white light (E), secondary minerals and kerogen in red (633 nm excitation; F), and filamentous kerogen structures in green (433 nm excitation; G) in another flame structure in the same thin section (Flame structure 3) where the filamentous nature of the kerogen structures is clearly visible.

In the high magnification Figs. 3E through 3G, the plain light image (Fig. 3E) shows bright-coloured refractive quartz crystals in white, while the dark central area contains kerogen and secondary minerals. The same area is shown via a 633 nm excitation wavelength (Fig. 3F), which causes both carbonaceous matter and distinct secondary crystals to auto-fluoresce. Notice the euhedral black spot in middle of the image, possibly the location of a zircon crystal. Fig. 3G shows the same area illuminated using a wavelength of 488 nm, where several linear features appear filamentous in nature in plan view. Also present are distinct bright globules, potentially kerogenous in nature, some appearing filamentous, and some bifurcation may be discernable. While some non-fluorescing grains can be seen by the near-total occlusion of the fluorescence signal (dark black areas), in general the structures fluorescing under 488 nm illumination do not appear to correspond for the most part to grain boundaries. The sample resolution is quite excellent, as details down to the 1 μm-scale are easily visible, the resolutions of which would only be improved with longer scan times (i.e. 8-10-12-hours vs. 6).

Fig. 4 shows the same thin section again, this time images are overlaid by image stacking to create a 1-D image, highlighting the difficulty in discerning features without 3-D imaging. This image is the same as the one above, but the rotation angle and 3-D nature allows for much better interpretation of the features observed. A conservative final analysis based on this 3-D image would be that the black areas represent regions of pure quartz that does not fluoresce under 488 or 633 nm excitations, while red areas represent secondary minerals (described above in the preliminary data) that produce an emission spectrum when exposed to 633 nm radiation. Unfortunately, this overlaps with kerogen excitation wavelengths and thus occludes much of the kerogenous matter we know is there (via Raman spectroscopy).

**Figure 4.**
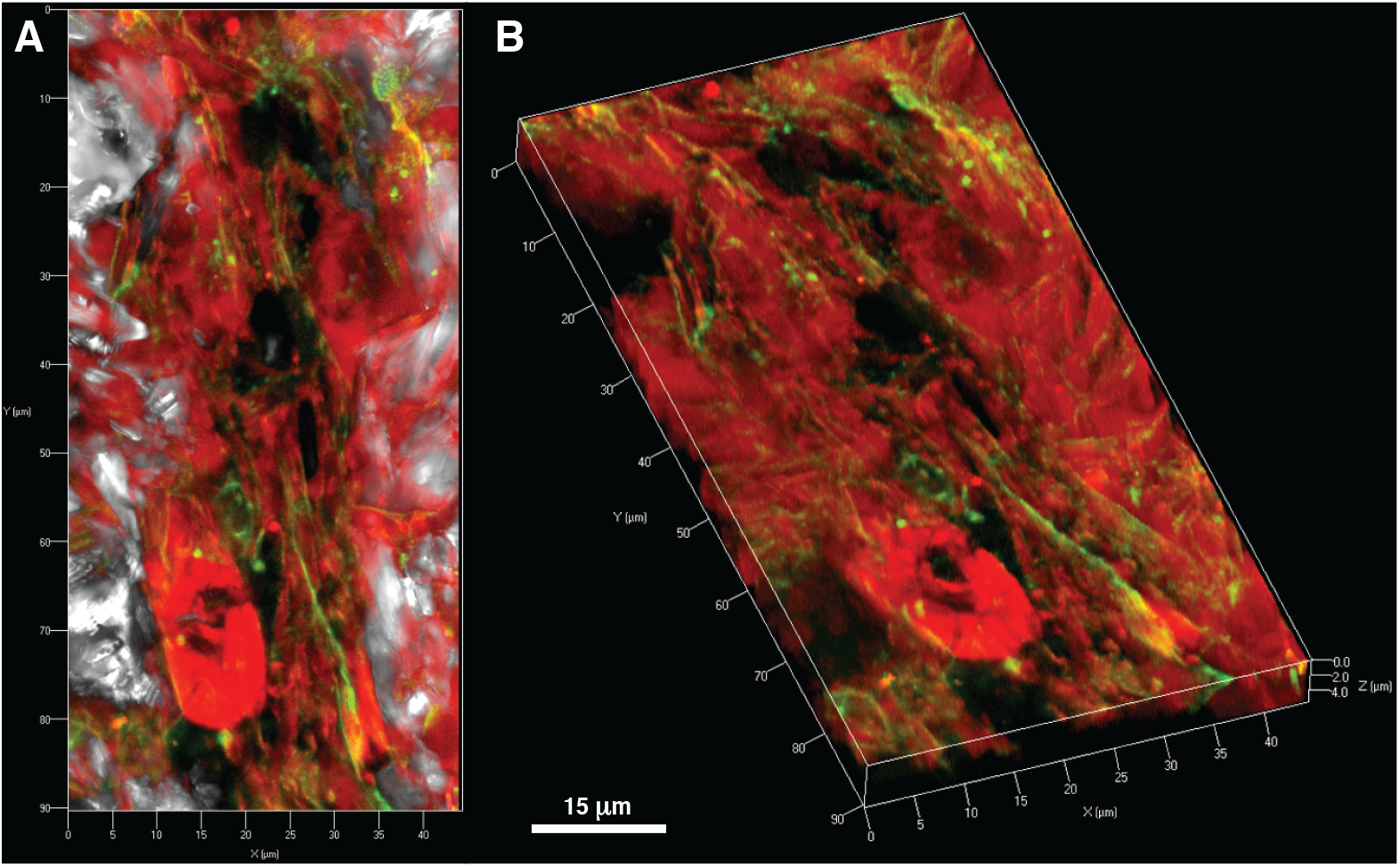
(A) Planar and (B) 3-dimensional z-stack imaging of thin section 11-452-1B5 showing different components (mats vs. quartz minerals vs. secondary minerals) of the Moodies mat occurrences. The wavelengths of excitation and their corresponding color channels are the same as represented in Figures 3E through 3G. This image highlights the added benefit of confocal imaging for confirming the nature of filamentous structures (e.g., sheet-like but linear in plan view vs. truly filamentous).

Images obtained using a green excitation wavelength (488 nm) reveal many linear features in line with the “up” direction of the flame structure. In most cases, it can be seen that the structures are nearly 1-dimensional, consistent with prior observations of filamentous microstructures that have been considered putative microfossils (e.g., Homann et al. 2015; 2018). However, the 3-D nature of the confocal image reveals that in some cases, what appear to be filamentous structures in plan view in fact extend into the sample, forming a 2-D sheet. Given that this is a flame structure, it is not unreasonable to conclude that the distortions and tendency for the green fluorescent matter to appear broken and/or to form a sheet like structure that follows grain boundaries is possibly the result of compression and fluid flow during the formation of the flame structure.

These observations are consistent with a biogenic origin modified by sedimentary processes, but like most microfossils that do not bear distinctively biological structures (e.g., septa visible in cells preserved during cell division; specific protruding or surface features that characterize test-forming eukaryotes, etc.), unfortunately positive identification with absolute certainty of these structures as fossilized cells is impossible. Nonetheless, the ensemble of evidence seen here is consistent with a biological origin.

In conclusion, the use of confocal imagery to determine the morphology of potentially biogenic structures is strongly affected by the duration and parameters of the scan, (longer, with more integration is better) laser wavelengths selected is critical, (i.e. interferences from other fluorescent minerals, vs. fluorescence from the kerogenous matter), the degree of alteration of the organic matter and potential growth of secondary minerals which may distort or destroy microbial morphologies, and finally the host medium of the potential fossils (the more homogeneous the host medium the better), as grain boundaries and reflection/refraction of grain edges at this scale become increasingly troublesome if we are searching for evidence of the most ancient forms of life (i.e. bacteria) which may only be 1-2 μm thick, akin to the distortions observed by primary and secondary grain boundaries and secondary crystal overgrowths.

### 4.2 Elemental and mineralogical evidence for primary vs. secondary processes via EMPA and DRX

BIF are chemical sedimentary deposits, primarily of Archean and Paleoproterozoic age, that constitute vast majority of Earth’s major economic iron ore deposits. They are also enigmatic in the mechanisms of their deposition, which are highly debated. The classic view holds that the oxidation of ferrous iron in surface waters resulted in the precipitation of amorphous iron (III) oxides (e.g., James, 1954) that are inherently unstable and underwent diagenetic and metamorphic transitions to more stable, dehydrated iron oxide minerals such as magnetite and hematite (see Konhauser et al. 2017 for review). Alternatively, it is also proposed that the precursor minerals were deposited as reduced ferrous iron minerals such as greenalite, minnesotaite, or siderite (e.g., Rasmussen et al. 2013), only to be oxidized during later fluid alteration. Furthermore, in the case of iron (III) oxide precursor minerals, it is thought that bacteria would have played a major role in their deposition (e.g., Konhauser et al. 2002), whether by oxygenic or anoxygenic Fe(II)-based photosynthesis. However, the mineralogy of BIF is in reality significantly more complicated than a simple mixture of iron (II) and (III) oxide minerals in a matrix of nanocrystalline quartz; they also often contain Fe-rich Ca and Mg carbonates (ankerite and dolomite), complex clay minerals such as stilpnomelane and annite, and trace rutile, apatite, and other accessory phases. These complex mineral assemblages may provide additional insight into depositional, diagenetic, and metamorphic conditions, and could also contribute to the debate surrounding the origins of these enigmatic deposits.

We analyzed selected excellently preserved drill core samples of the Dales Gorge member by μXRF, X-ray diffraction (XRD), and electron microprobe (EMPA) in order to better understand the mineralogical composition at fine scales and potential paleoenvironmental implications. Whole-rock powder analyses by μXRF confirms that the sample set, selected for its mineralogical diversity, spans a wide range in Fe and Si concentrations, accessory chemical sediment content (notably carbonate), and detrital sediment contributions (as reflected by the concentrations of detrital indicators Al and Ti) (Table 1). We chose two samples of contrasting overall composition for further analyses: DD98-30A, which is an iron-rich chemical sediment composed almost entirely of magnetite, hematite, quartz, and minor clays, and DD98-26A, a more cherty sample with more complex mineralogy composed of quartz, magnetite, dolomite, and higher concentrations of accessory clay minerals (Fig. 5).

**Table 1.** Major element composition of Dales Gorge Iron formation samples as determined by μXRF. Values are in weight percent.

|  | SiO<sub>2</sub> | Al<sub>2</sub>O<sub>3</sub> | TiO<sub>2</sub> | CaO | Na<sub>2</sub>O | K<sub>2</sub>O | MgO | Fe<sub>2</sub>O<sub>3</sub> | MnO | Total |
| --- | --- | --- | --- | --- | --- | --- | --- | --- | --- | --- |
| **DD98-1D** | 29.23 | 0.45 | 0.01 | 0.04 | 1.86 | 0.43 | 2.08 | 65.91 | 0.00 | 100.00 |
| **DD98-23A** | 52.60 | 1.28 | 0.04 | 3.38 | 0.05 | 1.15 | 2.84 | 38.34 | 0.32 | 100.00 |
| **DD98-23B** | 96.90 | 0.19 | 0.02 | 0.02 | 0.00 | 0.28 | 0.10 | 2.47 | 0.02 | 100.00 |
| **DD98-29A** | 58.98 | 0.00 | 0.01 | 1.54 | 0.98 | 0.08 | 1.48 | 36.92 | 0.02 | 100.00 |
| **DD98-29B** | 40.52 | 0.04 | 0.00 | 0.64 | 0.51 | 0.12 | 0.85 | 57.32 | 0.00 | 100.00 |

*Values in weight percent*

**Figure 5.**
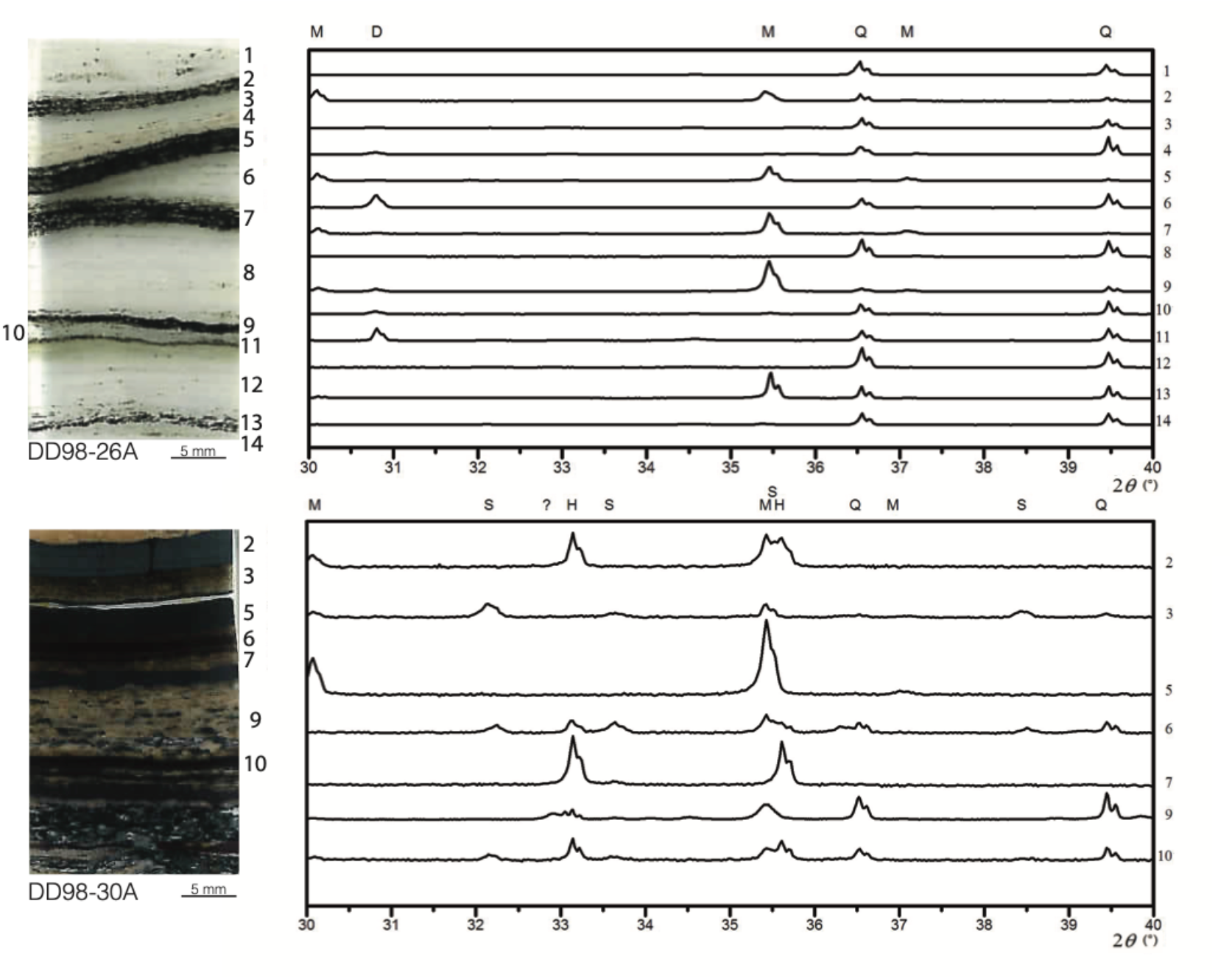
X-ray diffractograms for Dales Gorge Member samples DD98-26A and DD98-30A from the 2495 – 2454 Myr Brockman Iron Formation, Hamersley Group (W. Australia). See Figure 1 for approximate stratigraphic heights. H: hematite; M: magnetite; S: siderite, Q: quartz; D: dolomite.

For DD98-26A, the iron oxide minerals are highly localized to dark layers in an otherwise white quartz-rich matrix (Fig. 5). Both dolomite and clay contents were variable between the layers, with no clear relation to magnetite or quartz content, minor apatite was detected in three layers, and hematite was below quantification limits (Table 2).

**Table 2.** Mineral abundances (in percent) determined layer-by-layer for Dales Gorge samples DD98-26A and DD98-30A. See Figure 5 for corresponding spectra and thin section photographs.

| Sample | Layer | Quartz | Magnetite | Hematite | Dolomite | Clay Minerals | Other |
| --- | --- | --- | --- | --- | --- | --- | --- |
| **DD98-26A** | 1 | 74 | - | - | - | 16 | - |
|  | 2 | 70 | 15 | - | 1 | 14 | - |
|  | 3 | 41 | - | - | - | 58 | 1 |
|  | 4 | 94 | - | - | 2 | 3 | - |
|  | 5 | 45 | 39 | - | 3 | 8 | 6 |
|  | 6 | 83 | - | - | - | 3 | 14 |
|  | 7 | 42 | 39 | - | 4 | 12 | 3 |
|  | 8 | 85 | - | - | 1 | - | - |
|  | 9 | 36 | 29 | - | 5 | 27 | - |
|  | 10 | 70 | - | - | 4 | 23 | - |
|  | 11 | 62 | - | - | 8 | 29 | - |
|  | 12 | 54 | - | - | - | 45 | - |
|  | 13 | 44 | 9 | - | - | 46 | - |
|  | 14 | 42 | - | - | - | 57 | - |
| **DD98-30A** | 2 | 6 | 46 | 49 | - | - | - |
|  | 3 | 59 | 8 | - | - | 18 | 5 |
|  | 5 | 8 | 92 | - | - | - | - |
|  | 6 | 65 | 6 | 5 | - | 19 | 4 |
|  | 7 | 13 |  | 77 | - | 11 | - |
|  | 9 | 88 | 2 | 1 | - | 9 | - |
|  | 10 | 72 | 4 | 10 | - | 10 | 3 |

For DD98-30A, iron oxides are represented by nearly pure hematite, pure magnetite, and mixtures thereof. Dolomite was not detected, siderite was present in several layers at low abundances (<5%), and clay concentrations were significantly lower than for DD98-26A. For the two most iron oxide-rich bands, clays were below quantification limits. In both samples Mn- and K- bearing phases were detected in the most clay-rich intervals, but our data do not permit us to resolve whether they represent zones of intensified metasomatism or if the Mn- and K-enrichments reflect primary compositional features.

We also examined grain sizes (mixed with lattice distortion effects) using the Scherrer method (Scherrer, 1918) as implemented by the HighScore software (Malvern Panalytical, Ltd, Malvern, UK). This method evaluates peak broadening compared with a standard analyzed under the same conditions to infer crystallite size. The results are presented graphically in Fig. 6 as box and whisker plots showing the maximum and minimum sizes for each mineral, which are also tabulated in Supplemental Table 1. The quartz-rich sample DD98-26A generally shows a significantly larger range in indicated grain sizes, with quartz crystallographic domains reaching up to 3 μm compared to 1 μm in DD98-30A. Both samples show quartz crystallographic domains reaching down to 100 nm or below. Magnetite crystallographic domains reached a similarly higher range in DD98-26A relative to DD98-30A (1000 nm vs. 500 nm), whereas hematite, which is rare in DD98-26A, is found at significantly smaller in crystal sizes relative to DD98-30A (400 nm vs. 1000 nm). Inferred clay mineral sizes ranged from 200 nm to 20 nm and are comparable between the two samples.

**Figure 6.**
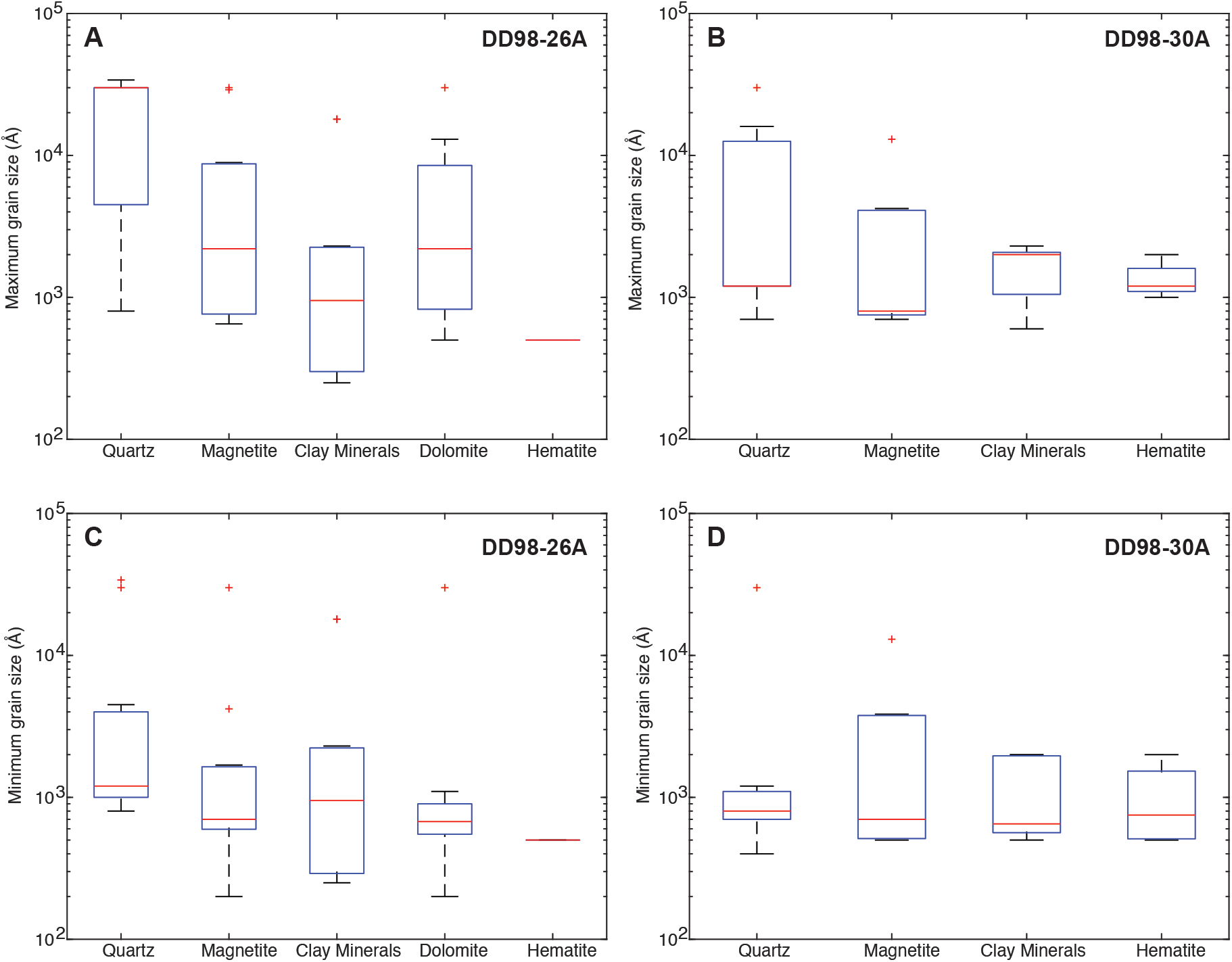
Box and whisker plots of grain size distributions in ångströms for the dominant minerals in Dales Gorge Member samples DD98-26A and DD98-30A. Maximum and minimum grain sizes are presented for sample DD98-26A in (A) and (C), respectively, and for DD98-30A in (B) and (D), respectively. Red lines represent the median of maximum or minimum values across the entire thin section, the box limits represent 25^th^ and 75^th^ percentiles, and whiskers represent the range of data not considered outliers (which appear as red crosses).

Electron microprobe elemental analyzer (EMPA) measurements and backscatter images reveal that these samples are much more complex at the micro-meter to nano-meter scale, and that bulk rock X-Ray diffraction studies represent a necessarily simplified picture. Similar to the XRD data, the EMPA analyses reveal a wide range in crystal sizes between different layers, even within a single thin section. For sample DD98-26A, Fig. 7 shows a progression in iron oxide crystal sizes (represented by the light-colored minerals in electron backscatter images) in the same thin section that in these images appears to correlate with iron mineral abundance. EMPA data for both iron oxides (in white) and quartz (in gray) reveals no clear trend in composition for iron-rich and silica-rich phases as a function of grain size, at least with respect to major elements (Table 3). Instead, the EMPA composition data strongly reflect the contrasting and bimodal mineralogy of chert-iron oxide couplets at all apparent grain sizes.

**Table 3.** Major element composition of Dales Gorge Iron formation samples as determined by EMPA (see Figure 7 for corresponding points). Values are in weight percent.

| Panel | Point | SiO<sub>2</sub> | Al<sub>2</sub>O<sub>3</sub> | CaO | Na<sub>2</sub>O | K<sub>2</sub>O | MnO | TiO<sub>2</sub> | P<sub>2</sub>O<sub>5</sub> | FeO | MgO | Total |
| --- | --- | --- | --- | --- | --- | --- | --- | --- | --- | --- | --- | --- |
| A | 1 | 42.52 | 2.04 | 0.02 | 0 | 3.99 | - | - | - | 34.96 | 5.79 | 100.67 |
|  | 2 | 0.11 | - | 26.94 | 0.07 | 0.01 | 1.09 | - | - | 17 | 8.51 | 53.81 |
| B | 3 | 96.2 | 0.03 | 0.01 | 0.23 | 0.01 | - | - | - | 1.21 | 0.22 | 97.96 |
|  | 4 | 97.83 | 0.01 | 0.01 | 0.02 | 0.03 | - | - | - | 0.89 | 0.15 | 98.94 |
|  | 5 | 91.35 | 0.02 | 0.04 | 0.02 | 0.03 | - | - | - | 0.04 | 0.02 | 91.54 |
|  | 6 | 97.09 | 0.05 | 0.02 | 0.04 | 0.1 | - | - | - | 3.01 | 0.35 | 100.67 |
| C | 7 | 101 | 0.06 | 0.03 | - | - | - | - | - | 1.06 | 0.01 | 102.17 |
|  | 8 | 0.83 | 0.01 | 0.02 | 0.002 | - | - | - | - | 92.14 | 0.04 | 93 |
|  | 9 | 0.06 | 0.03 | 27.08 | 0 | 0.002 | 0.95 | - | - | 18.73 | 7.92 | 54.87 |
| D | 10 | 0.83 | 0.03 | 0.04 | 0.01 | 0.01 | - | - | - | 92.42 | 0.01 | 93.4 |
|  | 11 | 1 | 0.03 | 26.49 | 0.01 | 0.01 | 0.74 | - | 0.42 | 18.47 | 8.18 | 56.05 |
|  | 12 | 99.96 | - | 0.03 | 0.01 | - | - | - | - | 0.06 | 0.03 | 100.67 |

MnO: aside from 3 analyses ranging from 0.74 to 1.09 percent, all other values are less than 0.01 percent. P<sub>2</sub>O<sub>5</sub>: with the exception of 1 analysis at 0.42 percent, all concentrations are below 0.03 percent. TiO<sub>2</sub> : all concentrations are largely below 0.01 percent

**Figure 7.**
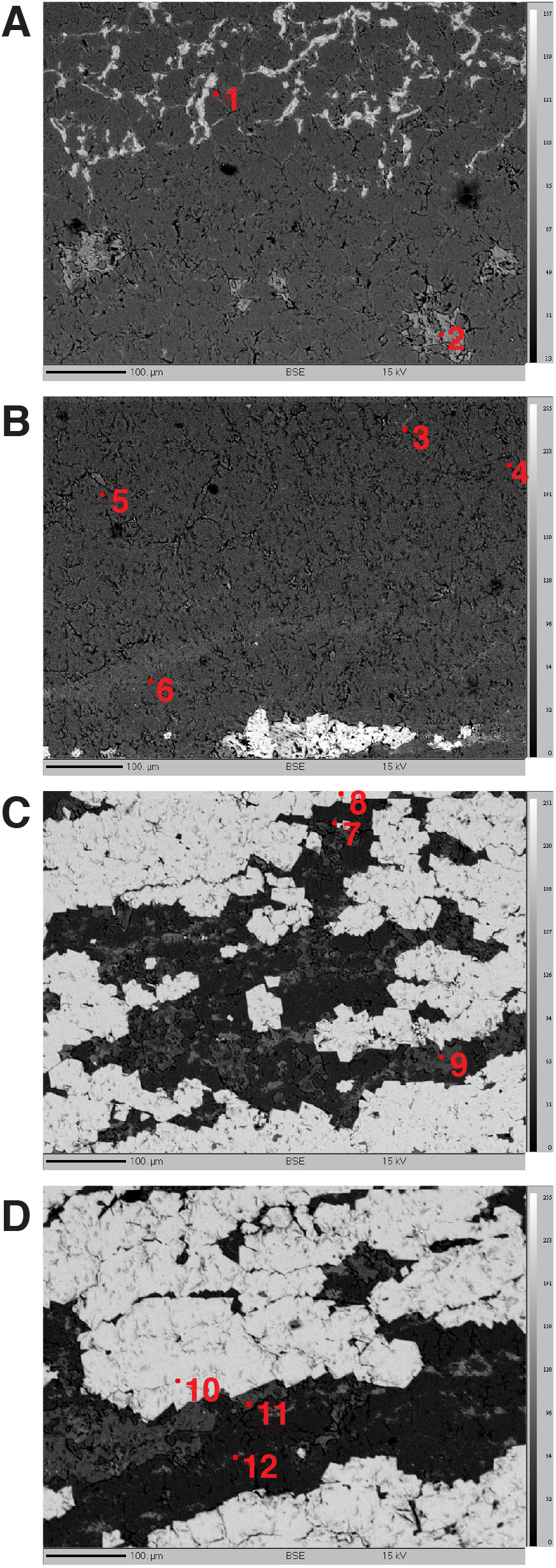
Electron Probe Microanalyzer (EPMA) backscatter images of thin section DD98-26A of the Dales Gorge Member of the Brockman iron formation. Major element data for the points of analyses are tabulated in Table 3.

As both samples come from the same drill core, it is unlikely that these differences in both mineralogy and crystal grain sizes may be related to differences in metamorphic conditions experienced by the two samples. Rather they more likely indicate primary compositional and local sedimentological and diagenetic control over the modal mineralogy of the samples, and in turn the control of the different mineralogical compositions over recrystallization processes that generated the grain size distributions observed. It is generally accepted that the primary mineralogy of BIF was originally hydrated amorphous mineral precipitates formed directly in the water column with nm-scale crystal domains that flocculated upon continued particle precipitation and aggregation (Konhauser et al. 2017).

The minerals deposited at the seafloor were thus meta-stable, and during early diagenesis and burial metamorphism they would have progressively dehydrated, continued to aggregate, and eventually undergone mineral transformation to more stable crystalline forms. The crystal domain size achieved would be subject to a complex combination of controls that would not only include temperature and pressure, but also pH, salinity, crystal growth rate, and the presence of organic or inorganic elements that may accelerate or poison crystal growth. Apparently, in the case of the contrasting iron-rich and silica-rich iron formation samples studied here, for which metamorphic conditions were effectively identical, it would appear that the more silica- and clay-rich sample, which presumably represents slower background deposition in the absence of strong iron resupply (Konhauser et al. 2017), achieved larger crystal sizes.

This is consistent with a sedimentation rate control over recrystallization, although compositional effects (e.g., the role of Fe doping in silica phases and vice-versa) should also be considered. This statistical and semi-quantitative examination of mineral sizes and distributions reveals previously unobserved trends between composition and crystal growth that has the potential to yield new insights into the paragenesis of BIF precursors from nm-scale amorphous flocculates to highly crystalline minerals stable in the rock record.

### 4.3 EPR as a proxy for evaluating the syngenicity

EPR is a non-destructive and non-invasive technique that is based on the adsorption of electromagnetic energy by paramagnetic nuclei and the re-emission of the incident electromagnetic energy at modified wavelength as the result of spin separation (the so-called Zeeman effect). It thus targets paramagnetic nuclei with unpaired free electrons. The exploited signal consists of a magnetic adsorption band whose shape and magnitude are related to the presence of specific paramagnetic nuclei, for example metals such as iron, but also radicals of non-metals such as carbon. The adsorption spectra may be fitted with different mathematical functions, such as Gaussian or Lorentzian equations, that account for the magnitude of the adsorption edge (A_pp_; Fig. 8).

**Figure 8.**
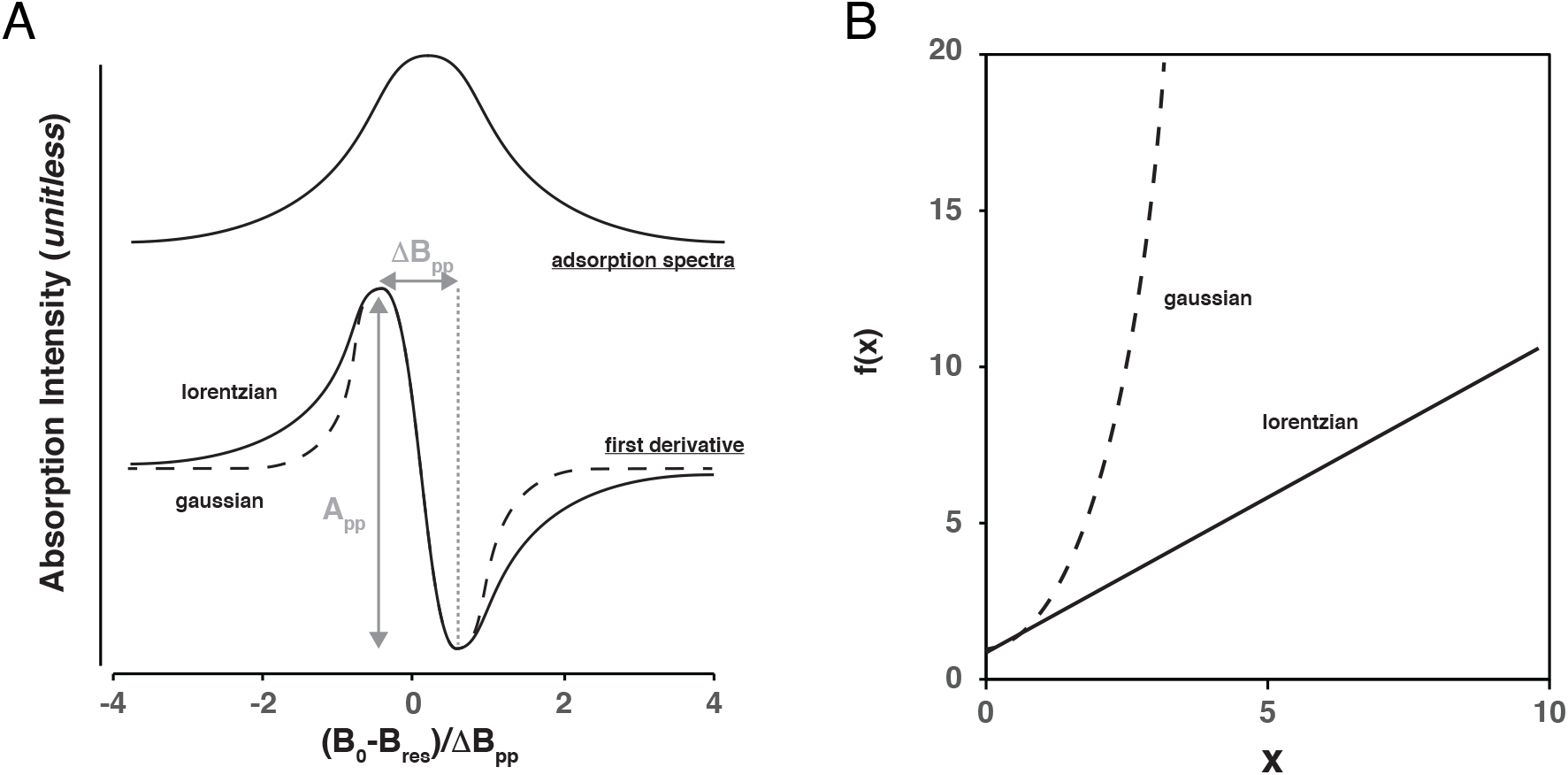
(A) Theoretical EPR spectra with gaussian (dashed line) and lorentzian (solid line) peak shapes. B_0_ refers to the ambient magnetic field, B_res_ is the magnetic field at the center of the resonance line, ΔB_pp_ represents the peak-to-peak width of magnetic resonance, and A_pp_ represents the amplitude of the resonance feature from peak to peak. (B) Representation gaussian and lorenztian fits in x-y space where x = ((B_0_ - B_res_) / ΔB_pp_)^2^ and f(x) is the square root of the product of ((B_0_ - B_res_) / ΔB_pp_) and A_pp_ / F(B_0_ – B_res_), where F = x + ¾ for a lorentzian fit and F=e^(x-1/4)^ for a gaussian fit (see Appendix in Bourbin et al. (2013) for a detailed description).

A Gaussian fit results from electron spin interacting with nuclear spin of neighbouring atoms (protons for example), or with neighbouring electron spin through dipolar magnetic interactions (Abragam, 1961). A Lorentzian fit results either from dipolar interactions in magnetically diluted systems or, on the contrary, in systems where the concentration of electron spin is high, the distances of interaction are reduced, and electron spins interact by exchange in the form of Coulomb interaction (Abragam, 1961).

In the specific case of carbon, it has been observed that young organic matter tends to show adsorption spectra that have an intermediate shape between a Gaussian and Lorentzian form, however with increasing age, tends towards a Lorentzien shape as the result of radiation-induced defects (Skrzypczak-Bonduelle et al. 2008; Bourbin et al. 2013).

This has led to the application of EPR analyses to the dating of ancient organic carbon in siliceous matrices. However, most analyses performed to date have focused on relatively pure cherts, despite the fact that cherty rocks may contain high concentrations of other elements, such as aluminum (in the case of cherty replacement of siliciclastic or volcanic rocks) or iron (in the case of Precambrian iron formations). We performed EPR analyses on powders from the Brockman iron formation to evaluate whether organic matter can be resolved by EPR in atypical cherty matrices and to evaluate EPR parameters commonly exploited for dating of organic matter by EPR. From these spectra, parameters such the value of the magnetic field at the center of the resonance line (B_res_), the peak-to-peak width of magnetic resonance (ΔB_pp_) and amplitude (A_pp_) (Fig. 8A) can be visually compared, and the form of the adsorption feature, specifically whether it is better described by a Gaussian, Lorentzian, or stretched Lorentzian function, can be evaluated (Fig. 8B). Departure of f(x) from a Lorentzian shape f_L_(x) is quantified by the correlation factor R10, considered an OM “internal clock” applicable over geological timescales (see Bourbin et al. 2013, for a detailed description):

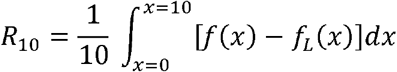

where *R*_10_ corresponds to the algebraic surface between the curve *f(x)* representing an experimental EPR spectrum and the curve *f_L_* representing a Lorentzian line. *R*10 is negative for a low-dimensional distribution (*D* <3) and positive for an EPR line intermediate between Lorentzian and Gaussian lines.

We see that R10 is a measure of the Lorentzian deviation because it is equal to zero when *f(x)* = *f_L_(x)*. On the other hand when R10 > 0, we have a close spin regime (quantum exchange regime) with a Gaussian / Lorentzian profile. Finally, when R10 < 0 we have a distant spin regime (dipolar regime) in a reduced space (D = 1, 2) with a stretched Lorentzian profile.

The method of calculating the factor R10 is very interesting because it makes it possible, from an EPR absorption spectrum, to extract the nature of the interactions between spins within the material (“quantum” [wave function] or “classical” [dipole], that is to say the interactions either “close” or “distant”). Moreover, it allows the determination of either “geometric” or “spatial” arrangement of the spins, for example in an “ordinary” 3-D arrangement, assembled in 2-D layers, or forming 1-D chains of the material. However, in this type of analysis, one should consider all the possible scenarios for both the exchange and the dipolar interaction (Tannous et Gieraltowski, 2021). It should be noted that Bourbin et al. (1993) and Skrzypczak-Bonduelle et al (2008) only considered the dipolar case at D = 1 and 2 without examining the exchange case.

Large-scale EPR scans (0 to 10,000 gauss) of DD98 samples reveal that they are similar in shape, but with contrasting intensities, regardless of temperature of analysis (Table 4; Fig. 9). The spectra are dominated by a strong positive peak below 1,000 gauss and a negative peak between 2,000 and 4,000 gauss. We attribute these features to the paramagnetic resonance of iron minerals, notably the Fe^3+^ dipolar interaction, and indeed, these spectra bear resemblance to similar scans of iron-rich potteries similarly dominated by the Fe^3+^ dipolar interaction (Watanabe et al. 2008).

**Table 4.** Maximum and minimum EPR intensities and corresponding field intensities for Dales Gorge BIF samples at two different temperatures (298° K, ambient temperature, and cooled to 150° K), as well as data for artificial mixtures of sample DD98-23A and a “pure” chert from the Buck Reef Chert, Onverwacht Group. Also shown are total iron concentrations (expressed as weight percent Fe_2_O_3_).

| Sample | Temperature | Fe<sub>2</sub>O<sub>3</sub> (wt. %) | SiO<sub>2</sub> (wt. %) | Intensity Max. | Intensity Min. | Field at Max (Gauss) | Field at Min (Gauss) |
| --- | --- | --- | --- | --- | --- | --- | --- |
| *Natural Samples, Dales Gorge BIF* |  |  |  |  |  |  |  |
| DD98-1D | 298K | 65.91 | 29.23 | 10190 | -12710 | 480 | 3050 |
| DD98-1D | 150K | 65.91 | 29.23 | 13800 | -15350 | 510 | 2820 |
| DD98-23A | 298K | 38.34 | 52.60 | 6300 | -7670 | 760 | 2830 |
| DD98-23A | 150K | 38.34 | 52.60 | 7340 | -7670 | 720 | 2430 |
| DD98-23B | 298K | 2.47 | 96.90 | 140 | 110 | 7530 | 3760 |
| DD98-23B | 150K | 2.47 | 96.90 | 140 | 110 | 7100 | 3410 |
| DD98-29A | 298K | 36.92 | 58.98 | 2290 | -2150 | 860 | 2450 |
| DD98-29A | 150K | 36.92 | 58.98 | 3080 | -2930 | 880 | 2390 |
| DD98-29B | 298K | 57.32 | 40.52 | 7140 | -10410 | 490 | 3540 |
| DD98-29B | 150K | 57.32 | 40.52 | 7450 | -9400 | 660 | 3240 |
| *Artificial mixtures between DD98-23A and "pure" chert from the Buck Reef Chert (Onverwacht Group)* |  |  |  |  |  |  |  |
| 25% Chert | 150K | 28.68 | 71.12 | 20637 | -19539 | 640 | 2260 |
| 50% Chert | 150K | 19.82 | 79.98 | 15015 | -13581 | 650 | 2050 |
| 75% Chert | 150K | 9.24 | 90.56 | 10715 | -9935 | 820 | 2150 |
| 95% Chert | 150K | 2.03 | 97.77 | 2100 | -1272 | 930 | 2100 |
| 100% Chert | 150K | 0.29 | 99.71 | 337 | -222 | 20 | 1630 |

**Figure 9.**
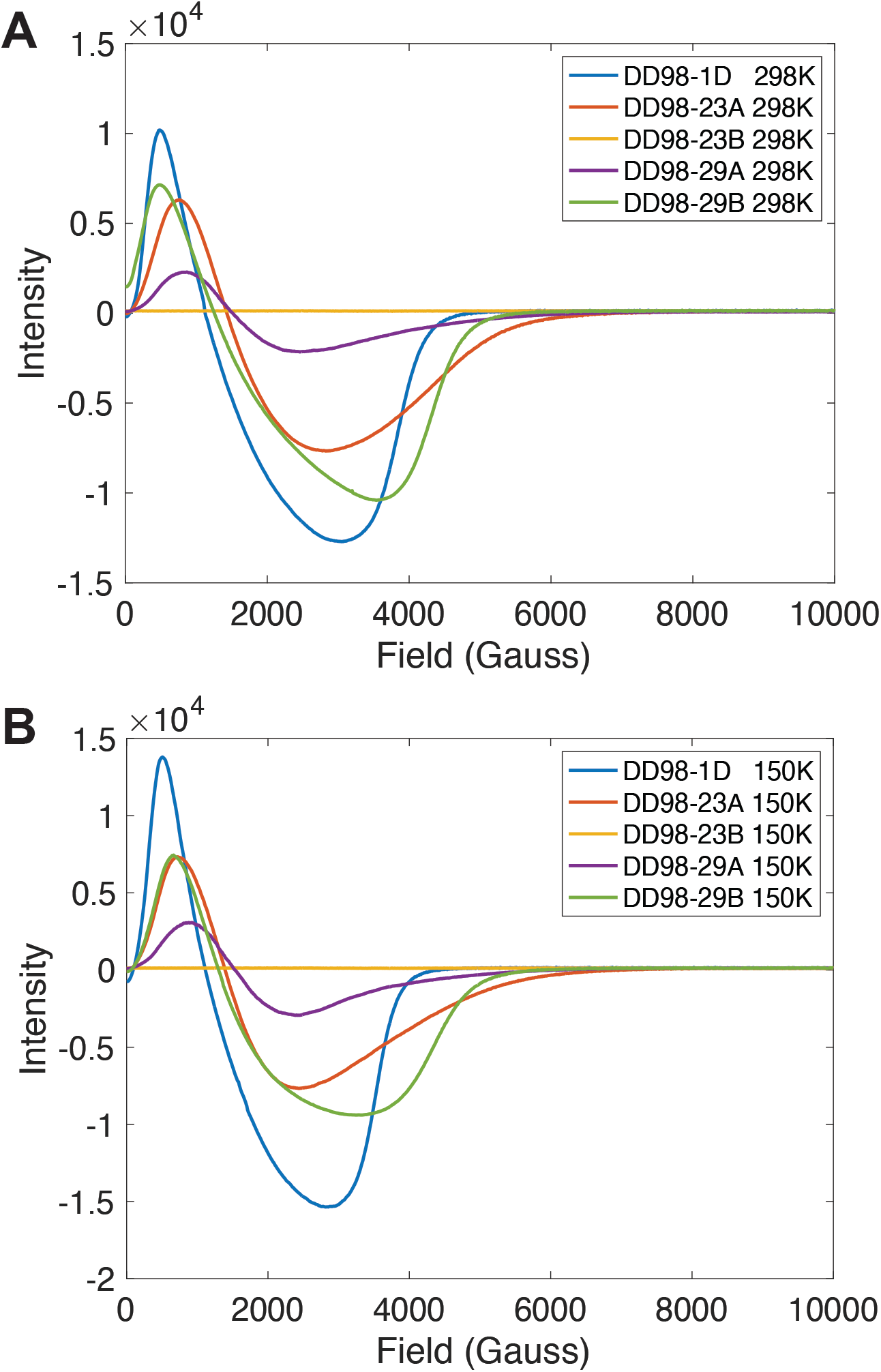
EPR spectra of selected Dales Gorge samples analyzed from zero to 10,000 Gauss (A) at room temperature and (B) at low temperature (150° K).

Fine-scale scans of these same samples over the resonance range expected for organic matter radicals revealed no resonance features, and thus no corresponding R10 factors could be determined. Surprisingly, this was the case even for the most iron-poor sample, which was composed of ~97 wt. percent SiO_2_; any signal from graphite was obscured by the Fe^3+^ dipolar interaction even at a total iron concentration of only 2.5 wt. percent (as Fe_2_O_3_). Iron is a common chert contaminant, and in to better understand its role in the detection and characterization of organic matter EPR, we performed artificial mixture experiments where we selected one iron-rich BIF sample (DD98-23A, 38.34 wt. percent Fe_2_O_3_) and mixed powders of this sample with those from a highly pure, organic-matter-rich black chert selected to represent a best-case scenario more comparable to the samples analyzed by Skrzypczak-Bonduelle et al. (2008) and Bourbin et al. (2013). This chert sample was collected from surface outcrop of the 3.42 Ga Buck Reef Chert of the Onverwacht Group, Barberton Greenstone Belt, S. Africa (see corresponding point on Figure 1). We can see that these artificial mixtures with different iron contents reproduce similar spectra to gross scans (Fig. 10) as the natural BIF samples presented in Fig. 9. While a clear resonance signal attributable to ancient organic matter was detected for the pure Buck Reef Chert end member, we see that even in the artificial mixture containing the least amount of iron formation powder, and only ~2% Fe_2_O_3_ by weight, resonance from organic matter radicals is wholly masked by the Fe^3+^ dipole interaction (Fig. 11).

**Figure 10.**
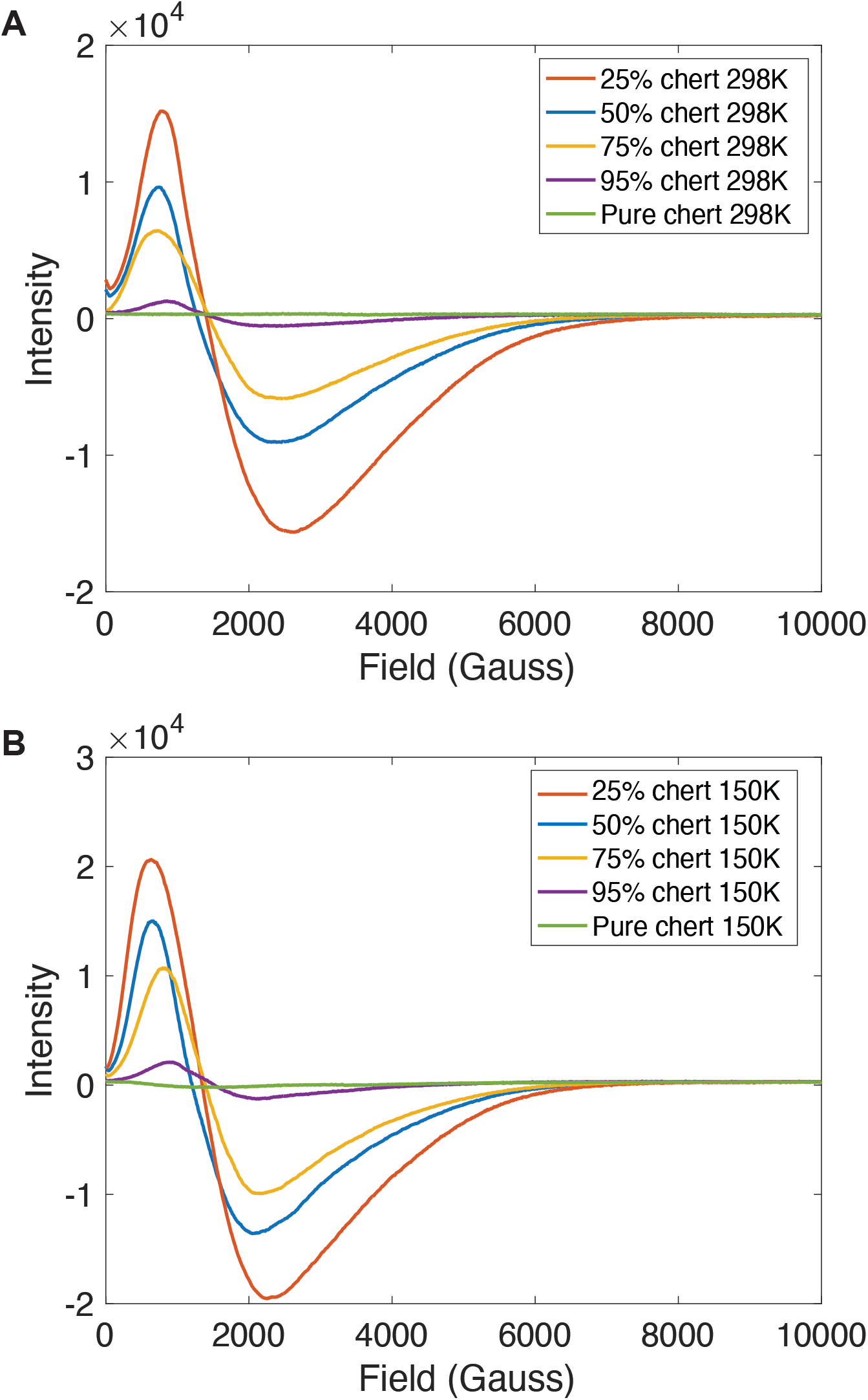
EPR spectra of black, organic-C-rich chert from the 3.42 Ga Buck Reef Chert, Barberton Greenstone Belt (“pure chert”) as well as artificial mixtures of chert and BIF powder (specifically, sample DD98-23A) analyzed from zero to 10,000 Gauss (A) at room temperature and (B) at low temperature (150° K). The artificial admixture of iron-rich siliceous material to chert reproduces the EPR intensity trends observed for the DD98 BIF sample set.

**Figure 11.**
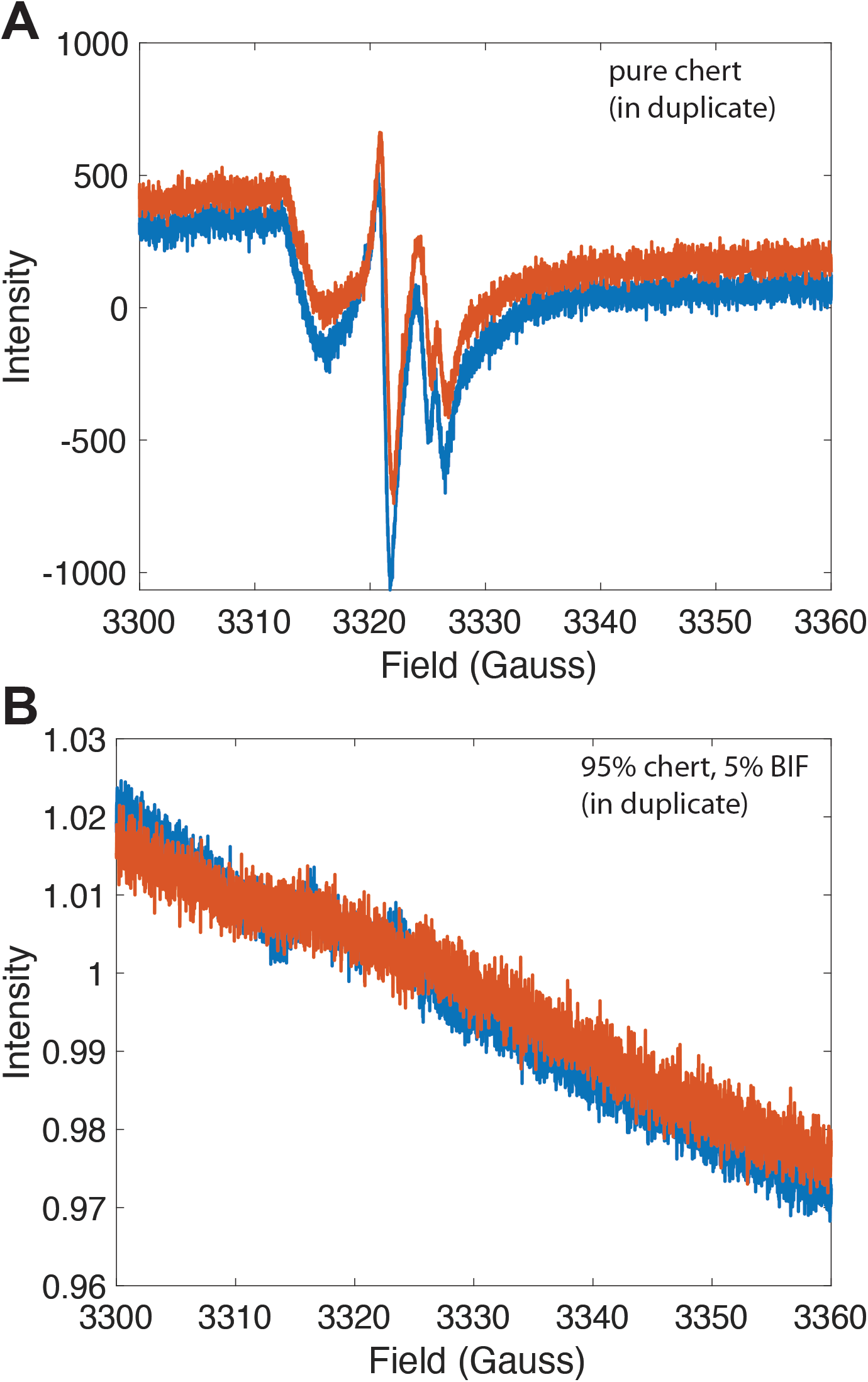
(A) High resolution spectra (analyzed in duplicate at 150° K) of organic radical EPR lines in black, organic-C-rich chert from the 3.42 Ga Buck Reef Chert, Barberton Greenstone Belt (“pure chert”). (B) The same sample to which 5% by mass of iron-rich BIF powder was added showing complete suppression of organic radical EPR lines. For B, spectra were normalized to the mean intensity of each spectrum.

While graphite was not detected by EPR in the natural BIF samples studied here, these new data do highlight the potential interest of obtaining chemically-pure kerogen via chemical extraction methods for Fe-enriched samples, or at least, of performing a careful selection of Fe-poor samples for EPR analysis (i.e. pure cherts), and that even very small residues of iron-bearing minerals may significantly affect RPE analyses. The effect of iron mineral content on RPE parameters, such as maximum and minimum intensity, is clearly visible in Fig. 12, where for both natural samples and artificial mixtures, maximum and minimum intensities broadly scale positively with Fe_2_O_3_ content, and inversely with SiO_2_ content, the two being related by their roles as the two major elements comprising typical BIF samples. It’s important to note that the relative proportions of iron oxide minerals that are ferrimagnetic (e.g., magnetite, hematite weakly at room temperature) or antiferromagnetic (e.g., hematite at low temperature) likely play an important role in determining the maximum and minimum intensities, and in this sample set, it would appear that magnetite content scales with total Fe oxide mineral content. Such correlation between electron spin parameters and Fe content has been observed for more recent chemical sediments (Crook et al. 2002). These results highlight the importance of controlling for the presence of additional paramagnetic substances during the analyses of residual organic matter in ancient chemical sediments, especially ancient ones that are likely to be rich in Fe-bearing minerals.

**Figure 12.**
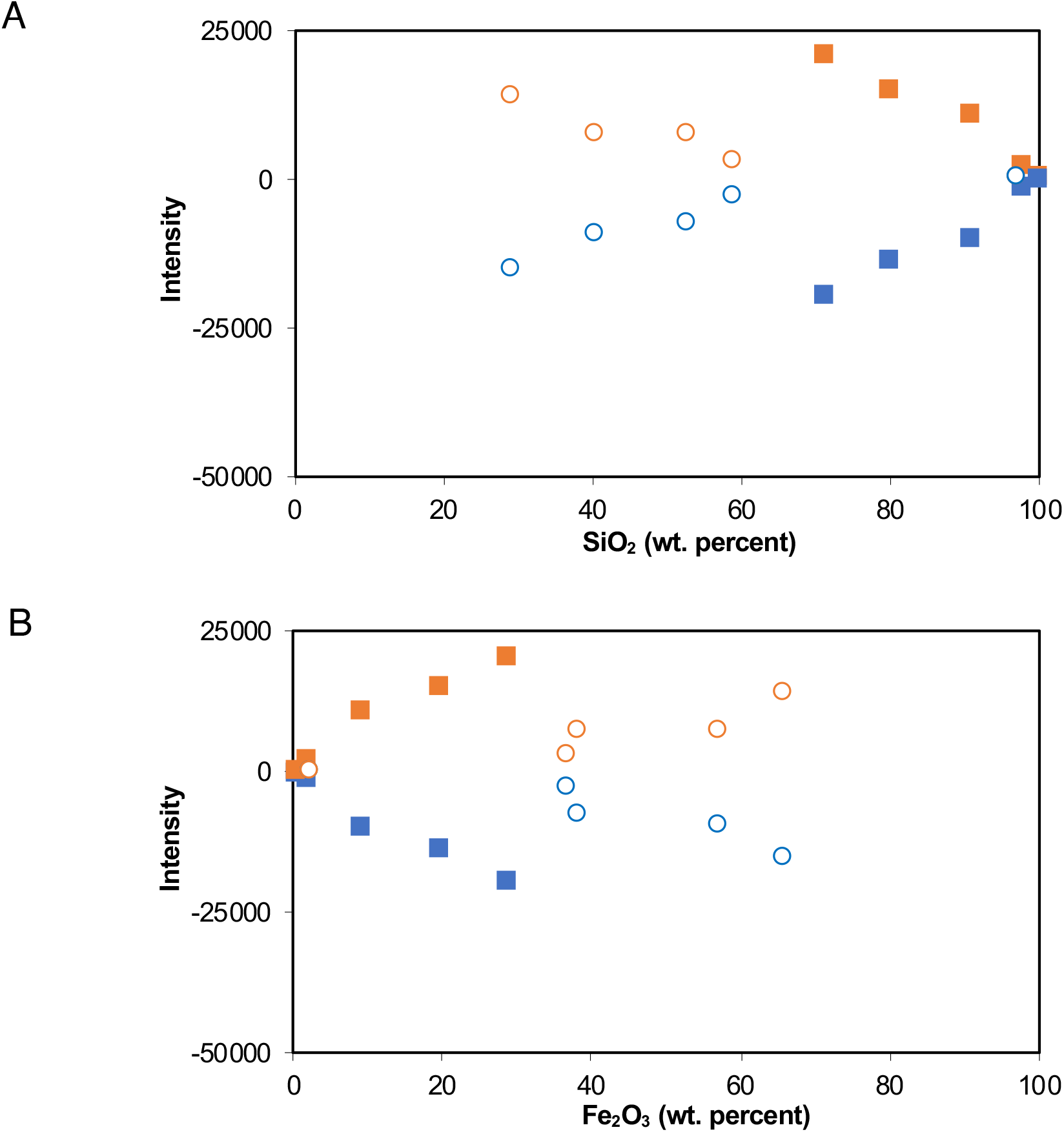
EPR maximum (orange) and minimum (blue) intensities as a function of Si (A) and Fe (B) concentrations for samples analyzed at low temperature (150° K). Open circles represent natural samples from the Brockman Iron Formation, filled squares represent artificial mixtures of sample DD98-23A (38.34% Fe_2_O_3_) and relatively pure natural chert from the 3.42 Ga Buck Reef Chert, Barberton Greenstone Belt (0.29% Fe_2_O_3_).

### 4.4 Synthesis

Claims of ancient biogenicity can be highly controversial even when multiple lines of evidence are presented (e.g., Dodd et al. 2017). In this study, we examined Archean and Paleoproterozoic sediments for which biological origins or strong biological influence is beyond dispute. Nonetheless, it can be seen here that the diverse techniques for evaluating biogenicity examined here, such as Confocal microscopy, Electron Microscopy, Electronic Paramagnetic Resonance analysis, and major element and mineralogical composition as determined by various X-ray and in-situ techniques, yield different types of information of varying utility for determining biogenicity. Direct imaging techniques such as confocal microscopy can be highly powerful tools for revealing biological structures and their physical relationships with host minerals, however they do not necessarily carry information regarding syngenicity where relationships with host minerals (e.g. overgrowths) or compaction features cannot be resolved. Techniques speaking more directly to degree of metamorphism and post-secondary modification, such as the analysis of secondary mineral assemblages by EMPA, μXRF, XRD, or EPR, may provide important information regarding the rock’s burial and metamorphic history, however this history may be difficult to simultaneously attribute to any biological structures or features that the sample may hold. EPR, which is relatively unique (along with Raman microscopy) in its ability to identify the degree of maturity of carbonaceous structures directly, can be hampered by the presence of additional paramagnetic phases, as demonstrated here.

Establishing biogenicity using micro- and nano-scale techniques clearly requires a multitude of evidence and corresponding techniques. Even then, traces of life in ancient rocks may be conflicting, even in the same singular sample. For example, in the case of the Moodies Group samples examined herein, we identified both putative ancient and possible modern microfossil structures. In the case of the Dales Gorge samples, the low preservation potential of organic carbon due to the iron-oxide rich composition of the samples, combined with pervasive recrystallization (even at low metamorphic grade and finely-sized crystallite formation), renders these putative “biological sediments” practically free of detectable biogenic structures.

In conclusion, traces of life may be difficult to detect with confidence even in the most biologically-influenced samples, despite a plethora of micro- to nano-scale techniques available for characterizing different potential biogenic aspects. At the very least, the work should help guide future researchers in their selection of techniques for establishing biogenicity, in full consideration of the advantages and pitfalls.

## 5. Conclusion

Micro- to nano-scale techniques in materials characterization are also powerful tools for the study of Earth’s primitive biosphere. In this study we applied confocal laser microscopy, visible light and electron microscope imaging, spatially-resolved X-Ray diffraction, electron microprobe elemental mapping, and electron paramagnetic resonance analysis to two interesting cases where biogenicity has been debated: filamentous kerogenous structures preserved sandstones of the 3.22 Ga Moodies Group (Barberton Greenstone Belt, S. Africa), and finely laminated oxide-facies BIF from the 2.48 Ga Dales Gorge Member of the Brockman Iron Formation (Hamersely Basin, W. Australia). In the case of the Moodies group filamentous structures, confocal laser microscopy clearly reveals the 3D nature of the filaments as well as the presence of metamorphic minerals cross cutting the filaments. These relationships strongly support a primary origin for the filamentous structures, consistent with the hypothesis that they represent syngenetic, fossilized filamentous bacteria. In the case of the Dales Gorge Member, μXRF, XRD, and EMPA analyses reveal large diversity in mineral abundance, size ranges, and composition, even at the individual layer and thin section scale, which, considering that these samples all experience similar metamorphic conditions, likely reflects complex and variable depositional and diagenetic conditions that exerted control over crystal size post-metamorphism. EPR analysis of trace kerogens in whole-rock powders of the Dales Gorge BIF samples reveals the overwhelming effect of magnetite ferromagnetism on the EPR spectra and suggests that extra caution is warranted during the analysis of ancient kerogens by EPR when magnetite or other iron (III) oxide minerals may be present. Collectively, this study highlights novel applications of micro- to nano-scale techniques of materials characterization as applied to putative primitive Earth fossils and their potential benefits for future paleobiological studies.

## Credit authorship contribution statement

MH, IF, and SL provided samples and assisted with *in-situ* characterization. FM performed XRD analyses and GS and SC performed EPR analyses. HB, JG, and SVL conceived the study and wrote the manuscript with contributions from all co-authors.

## Acknowledgments

We thank Jean-Pierre Oldra for thin sections, Jessica Langlade at Microprobe and Philippe Elies from UBO Plateforme d’Imagerie et de Mesures en Microscopie, all for assistance with sample preparation and analyses. We also thank Marcel Koken for stimulating discussions, and Charbel Tannous for his careful mathematical analysis of RPE curves. We are grateful to IUEM for providing financial support with UBO funds. Finally, we also thank Bertrand Sichler, who was among the first participants in the UBO Nano Research group, and critically read the text.

## References

Abragam, A. Principles of Nuclear Magnetic Resonance, Oxford, New York, (1961)

Bourbin, M., Gourier, D., Derenne, S., Binet, L., Le Du, Y., Westall, F., Kremer, B., Gautret, P., 2013. Dating Carbonaceous Matter in Archean Cherts by Electron Paramagnetic Resonance. Astrobiology 13, 151–162. doi:10.1089/ast.2012.0855

Brasier, M.D., Green, O.R., Jephcoat, A.P., Kleppe, A.K., Van Kranendonk, M.J., Lindsay, J.F., Steele, A., Grassineau, N.V., 2002. Questioning the evidence for Earth’s oldest fossils. Nature 416, 76–81. doi:10.1038/416076a

Crook, N.P., Hoon, S.R., Taylor, K.G., Perry, C.T., 2002. Electron spin resonance as a high sensitivity technique for environmental magnetism: determination of contamination in carbonate sediments. Geophys J Int 149, 328–337. doi:10.1046/j.1365-246X.2002.01647.x

de Ronde, C.E.J., Kamo, S.L., 2000. An Archaean arc-arc collisional event: a short-lived (ca 3 Myr) episode, Weltevreden area, Barberton greenstone belt, South Africa. Journal of African Earth Sciences 30, 219–248. doi:10.1016/S0899-5362(00)00017-8

Dodd, M.S., Papineau, D., Grenne, T., Slack, J.F., Rittner, M., Pirajno, F., O’Neil, J., Little, C.T.S., 2017. Evidence for early life in Earth’s oldest hydrothermal vent precipitates. Nature Publishing Group 543, 60–64. doi:10.1038/nature21377

Fel’dman EB, Lacelle S: Configurational averaging of dipolar interactions in magnetically diluted spin networks. J. Chem. Phys., 104: 2000–2010 (1996).

Gourier, D., Binet, L., Calligaro, T., Cappelli, S., Vezin, H., Bréhéret, J., Hickman-Lewis, K., Gautret, P., Foucher, F., Campbell, K., Westall, F., 2019. Extraterrestrial organic matter preserved in 3.33 Ga sediments from Barberton, South Africa. Geochimica et Cosmochimica Acta 258, 207–225. doi:10.1016/j.gca.2019.05.009

Gross, G.A., 1965. Geology of iron deposits in Canada. I. General geology and evaluation of iron deposits, Geol. Surv. Can., Econ. Geol. Rep. 22. doi:10.4095/102455

Heubeck, C., 2009. An early ecosystem of Archean tidal microbial mats (Moodies Group, South Africa, ca. 3.2 Ga). Geology 37, 931–934. doi:10.1130/G30101A.1

Heubeck, C., Engelhardt, J., Byerly, G.R., Zeh, A., Sell, B., Luber, T., Lowe, D.R., 2013. Timing of deposition and deformation of the Moodies Group (Barberton Greenstone Belt, South Africa): Very-high-resolution of Archaean surface processes. Precambrian Research. doi:10.1016/j.precamres.2013.03.021

Homann, M., Heubeck, C., Airo, A., Tice, M.M., 2015. Morphological adaptations of 3.22 Ga-old tufted microbial mats to Archean coastal habitats (Moodies Group, Barberton Greenstone Belt, South Africa). Precambrian Research 266, 47–64. doi:10.1016/j.precamres.2015.04.018

Homann, M., Sansjofre, P., van Zuilen, M., Heubeck, C., Gong, J., Killingsworth, B., Foster, I.S., Airo, A., Van Kranendonk, M.J., Ader, M., Lalonde, S.V., 2018. Microbial life and biogeochemical cycling on land 3,220 million years ago. Nature Geoscience 11, 1–8. doi:10.1038/s41561-018-0190-9

James, H.L., 1954. Sedimentary facies of iron-formation. Economic Geology 49, 235–293.

Johannessen, K.C., Vander Roost, J., Dahle, H., Dundas, S.H., Pedersen, R.B., Thorseth, I.H., 2017. Environmental controls on biomineralization and Fe-mound formation in a low-temperature hydrothermal system at the Jan Mayen Vent Fields. Geochimica et Cosmochimica Acta 202, 101–123. doi:10.1016/j.gca.2016.12.016

Karl, D.M., Brittain, A.M., Tilbrook, B.D., 1989. Hydrothermal and microbial processes at Loihi Seamount, a mid-plate hot-spot volcano. Deep-sea research. Part A. Oceanographic research papers 36, 1655–1673. doi:10.1016/0198-0149(89)90065-4

Konhauser, K.O., Hamade, T., Raiswell, R., Morris, R.C., Ferris, F.G., Southam, G., Canfield, D.E., 2002. Could bacteria have formed the Precambrian banded iron formations? Geology 30, 1079. doi:10.1130/0091-7613(2002)030<1079:CBHFTP>2.0.CO;2

Konhauser, K.O., Planavsky, N.J., Hardisty, D.S., Robbins, L.J., Warchola, T.J., Haugaard, R., Lalonde, S.V., Partin, C.A., Oonk, P.B.H., Tsikos, H., Lyons, T.W., Bekker, A., Johnson, C.M., 2017. Iron formations: A global record of Neoarchaean to Palaeoproterozoic environmental history. Earth Science Reviews 172, 140–177. doi:10.1016/j.earscirev.2017.06.012

Krapez, B., Barley, M.E., Pickard, A.L., 2003. Hydrothermal and resedimented origins of the precursor sediments to banded iron formation: sedimentological evidence from the Early Palaeoproterozoic Brockman Supersequence of Western Australia. Sedimentology 50, 979–1011. doi:10.1046/j.1365-3091.2003.00594.x

Lowe, D.R., Byerly, G.R., 1999. Geologic Evolution of the Barberton Greenstone Belt, South Africa. Geological Society of America. doi:10.1130/SPE329

Miyano, T., Miyano, S., 1982. Ferri-annite from the Dales Gorge Member iron-formations, Wittenoom area, Western Australia. American Mineralogist 1–16.

Noffke, N., Eriksson, K.A., Hazen, R.M., Simpson, E.L., 2006. A new window into Early Archean life: Microbial mats in Earth’s oldest siliciclastic tidal deposits (3.2 Ga Moodies Group, South Africa). Geology 34, 253. doi:10.1130/G22246.1

Pickard, A., 2002. SHRIMP U–Pb zircon ages of tuffaceous mudrocks in the Brockman Iron Formation of the Hamersley Range, Western Australia. Australian Journal of Earth Sciences 49, 491–507. doi:10.1016/0006-2944(75)90147-7

Rasmussen, B., Meier, D.B., Krapež, B., Muhling, J.R., 2013. Iron silicate microgranules as precursor sediments to 2.5-billion-year-old banded iron formations. Geology 41, 435–438. doi:10.1130/G33828.1

Scherrer, P., 1918. Bestimmung der Grösse und der inneren Struktur von Kolloidteilchen mittels Röntgensrahlen [Determination of the size and internal structure of colloidal particles using X-rays]. Nachr. Ges. Wiss. Goettingen, Math-Phys Kl 98–100.

Schopf, J.W., 1993. Microfossils of the early Archean apex chert: New evidence of the antiquity of life 260, 640–646. doi:10.1126/science.260.5108.640

Shapiro, R.S., Konhauser, K.O., 2015. Hematite-coated microfossils: primary ecological fingerprint or taphonomic oddity of the Paleoproterozoic? Geobiology 13, n/a–n/a. doi:10.1111/gbi.12127

Skrzypczak-Bonduelle, A., Binet, L., Delpoux, O., Vezin, H., Derenne, S., Robert, F., Gourier, D., 2008. EPR of Radicals in Primitive Organic Matter: A Tool for the Search of Biosignatures of the Most Ancient Traces of Life. Applied Magnetic Resonance 33, 371–397. doi:10.1007/s00723-008-0083-y

Tannous, C., Gieraltowski, J., 2021. Information theory based Electron Paramagnetic Resonance dating. ArXiv pre-print. https://arxiv.org/pdf/2105.01971.pdf

Tice, M.M., Bostick, B.C., Lowe, D.R., 2004. Thermal history of the 3.5–3.2 Ga Onverwacht and Fig Tree Groups, Barberton greenstone belt, South Africa, inferred by Raman microspectroscopy of carbonaceous material. Geology 32, 37–4. doi:10.1130/G19915.1

Trendall, A., Compston, W., de LAETER, D., Bennett, V., 2004. SHRIMP zircon ages constraining the depositional chronology of the Hamersley Group, Western Australia. Australian Journal of Earth Sciences 51, 621–644. doi:10.1111/j.1400-0952.2004.01082.x

Trendall, A.F., 1966. Altered pyroclastic rocks in iron-formation in the Hamersley range, Western Australia [discussion]. Economic Geology 61, 1451–1458. doi:10.2113/gsecongeo.61.8.1451

Trendall, A.F., Blockley, J.G., 1970. The Iron Formations of The Precambrian Hamersley Group Western Australia With Special Reference to the Associated Crocidolite. Geological Survey of Western Australia Bulletin 119, 1–366.

Watanabe, S., Farias, T.M.B., Gennari, R.F., Ferraz, G.M., Kunzli, R., Chubaci, J.F.D., 2008. Chemical process to separate iron oxides particles in pottery sample for EPR dating. Spectrochimica Acta - Part A: Molecular and Biomolecular Spectroscopy 71, 1261–1265. doi:10.1016/j.saa.2008.03.034

Xie, X., Byerly, G.R., Ferrell, R.E., Jr, 1997. IIb trioctahedral chlorite from the Barberton greenstone belt: crystal structure and rock composition constraints with implications to geothermometry. Contributions to Mineralogy and Petrology 126, 275–291. doi:10.1007/s004100050250

