## Supplemental Table 1 for "Micro- to nano-scale investigation of Precambrian metasediments: biogenicity and preservation in the 3.22 Ga Moodies Group (Barberton Greenstone Belt, S. Africa) and the 2.46 Ga Brockman Iron Formation (Hamersley Basin, W. Australia)"

| Sample | Mineral | Zone in Figure 5 | Max size (Å) | Min size (Å) |
|---|---|---|---|---|
| DD98-26A | Quartz | 1 | 34000 | 800 |
| DD98-26A | Clay | 1 | 300 | 300 |
| DD98-26A | Magnetite | 1 | 650 | 650 |
| DD98-26A | Quartz | 2 | 34000 | 800 |
| DD98-26A | Clay | 2 | 800 | 800 |
| DD98-26A | Magnetite | 2 | 2200 | 700 |
| DD98-26A | Hematite | 2 | 500 | 500 |
| DD98-26A | Quartz | 3 | 6100 | 1200 |
| DD98-26A | Clay | 3 | 1100 | 1100 |
| DD98-26A | Quartz | 4 | 32000 | 1000 |
| DD98-26A | Clay | 4 | 1100 | 1100 |
| DD98-26A | Magnetite | 4 | 800 | 200 |
| DD98-26A | Dolomite | 4 | 500 | 200 |
| DD98-26A | Quartz | 5 | 30000 | 1500 |
| DD98-26A | Clay | 5 | 18000 | 18000 |
| DD98-26A | Magnetite | 5 | 29000 | 4200 |
| DD98-26A | Dolomite | 5 | 30000 | 30000 |
| DD98-26A | Quartz | 6 | 800 | 34000 |
| DD98-26A | Clay | 6 | 2300 | 2300 |
| DD98-26A | Dolomite | 6 | 4000 | 500 |
| DD98-26A | Quartz | 7 | 2300 | 1200 |
| DD98-26A | Clay | 7 | 300 | 300 |
| DD98-26A | Magnetite | 7 | 2200 | 850 |
| DD98-26A | Dolomite | 7 | 850 | 650 |
| DD98-26A | Quartz | 8 | 30000 | 1000 |
| DD98-26A | Dolomite | 8 | 2200 | 1100 |
| DD98-26A | Quartz | 9 | 30000 | 1000 |
| DD98-26A | Clay | 9 | 18000 | 18000 |
| DD98-26A | Magnetite | 9 | 2200 | 700 |
| DD98-26A | Dolomite | 9 | 13000 | 700 |
| DD98-26A | Quartz | 10 | 4000 | 1000 |
| DD98-26A | Clay | 10 | 500 | 500 |
| DD98-26A | Magnetite | 10 | 1100 | 500 |
| DD98-26A | Dolomite | 10 | 2200 | 600 |
| DD98-26A | Quartz | 11 | 4500 | 4500 |
| DD98-26A | Clay | 11 | 250 | 250 |
| DD98-26A | Magnetite | 11 | 650 | 650 |
| DD98-26A | Dolomite | 11 | 800 | 700 |
| DD98-26A | Quartz | 12 | 6000 | 4000 |
| DD98-26A | Quartz | 13 | 30000 | 30000 |
| DD98-26A | Magnetite | 13 | 30000 | 30000 |
| DD98-26A | Quartz | 14 | 30000 | 4000 |
| DD98-30A | Quartz | 2 | 1200 | 400 |
| DD98-30A | Magnetite | 2 | 800 | 500 |
| DD98-30A | Hematite | 2 | 1200 | 500 |
| DD98-30A | Quartz | 3 | 1200 | 700 |
| DD98-30A | Clay | 3 | 1200 | 500 |
| DD98-30A | Magnetite | 3 | 800 | 800 |
| DD98-30A | Quartz | 5 | 700 | 700 |
| DD98-30A | Magnetite | 5 | 1300 | 500 |
| DD98-30A | Quartz | 6 | 2300 | 800 |
| DD98-30A | Clay | 6 | 2300 | 650 |
| DD98-30A | Magnetite | 6 | 13000 | 13000 |
| DD98-30A | Hematite | 6 | 1200 | 500 |
| DD98-30A | Quartz | 7 | 1200 | 1200 |
| DD98-30A | Clay | 7 | 600 | 600 |
| DD98-30A | Hematite | 7 | 2000 | 2000 |
| DD98-30A | Quartz | 9 | 30000 | 30000 |
| DD98-30A | Clay | 9 | 2000 | 2000 |
| DD98-30A | Quartz | 10 | 16000 | 800 |
| DD98-30A | Magnetite | 10 | 700 | 700 |
| DD98-30A | Clay | 10 | 2000 | 2000 |
| DD98-30A | Hematite | 10 | 1000 | 1000 |

**Supplemental Table 1.** Inferred maximum and minimum grain sizes for minerals identified by XRD in Dales Gorge samples.
